# A species-specific retrotransposon drives a conserved *Cdk2ap1* isoform essential for preimplantation development

**DOI:** 10.1101/2021.03.24.436683

**Authors:** Andrew Modzelewski, Wanqing Shao, Jingqi Chen, Angus Lee, Xin Qi, Mackenzie Noon, Kristy Tjokro, Gabriele Sales, Anne Biton, Terence Speed, Zhenyu Xuan, Ting Wang, Davide Risso, Lin He

## Abstract

Retrotransposons mediate gene regulation in multiple developmental and pathological processes. Here, we characterized the transient retrotransposon induction in preimplantation development of eight mammalian species. While species-specific in sequences, induced retrotransposons exhibit a similar preimplantation profile, conferring gene regulatory activities particularly through LTR retrotransposon promoters. We investigated a mouse-specific MT2B2 retrotransposon promoter, which generates an N-terminally truncated, preimplantation-specific *Cdk2ap1*^*ΔN*^ isoform to promote cell proliferation. *Cdk2ap1*^*ΔN*^ functionally contrasts to the canonical *Cdk2ap1*, which represses cell proliferation and peaks in mid-gestation stage. The mouse-specific MT2B2 element is developmentally essential, as its deletion abolishes *Cdk2ap1*^*ΔN*^, reduces cell proliferation and impairs embryo implantation. Intriguingly, Cdk2ap1^*Δ*N^ is evolutionarily conserved across mammals, driven by species-specific promoters. The distinct preimplantation Cdk2ap1^*Δ*N^ expression across different mammalian species correlates with their different duration in preimplantation development. Hence, species-specific transposon promoters can yield evolutionarily conserved, alternative protein isoforms, bestowing them with new functions and species-specific expression to govern essential biological divergence.

**One Sentence Summary:** In mammalian preimplantation embryos, retrotransposon promoters generate conserved gene isoforms, confer species-specific expression, and perform essential developmental functions.

## Main Text

Transposable elements constitute approximately 40% of mammalian genomes, underscoring their efficacy in exploiting host machinery for widespread propagation(*1, 2*). The mammalian mobilome is largely derived from three classes of retrotransposons; Long Terminal Repeat (LTR) retrotransposons, Long Interspersed Nuclear Elements (LINEs) and Short Interspersed Nuclear Elements (SINEs), all of which amplify via RNA intermediates using a “copy and paste” mechanism(*3, 4*). Once regarded as parasitic or “junk” DNA, emerging evidence suggests that specific retrotransposons are integral functional components of their host genome(*5*–*7*). Viral proteins encoded by certain retrotransposons have been co-opted by hosts for important developmental functions, such as placental cytotrophoblast fusion in mammals, telomere maintenance in *Drosophila*, and intracellular RNA transport across neurons (*8*–*13*). Yet the more prevalent retrotransposon exaptation is providing cis-regulatory elements that reprogram host gene expression in various developmental and pathological processes (*14*–*22*). Originating from ancient, exogenous retroviruses that no longer mobilize, a subset remain largely intact with LTR elements still harboring functional proviral sequences with intrinsic promoter activity, enhancer activity and splicing donor/acceptor sequences (*16, 23*–*29*), which substantially expand host gene regulatory landscape and transcript diversity. Due to their unique evolutionary history, retrotransposon mediated gene regulation is often species-specific(*25*), and its functional importance *in vivo* remains largely unclear.

Most retrotransposon integrations are thought to be deleterious to genome integrity, necessitating inactivation through degenerative mutations or epigenetic silencing during most developmental processes (*30*). However, potent retrotransposon induction occurs under specific developmental, physiological and pathological contexts, including preimplantation development(*31*–*33*), germ cell development(*34*–*36*), immune response(*37*–*39*), aging(*40*–*42*) and cancer(*28, 37, 43, 44*). In particular, a hallmark of mammalian preimplantation embryos is transient and robust retrotransposon induction, presumably the result of extensive epigenetic remodeling at the onset of early cell fate specification(*45*).

To comprehensively profile the retrotransposon landscape in mammalian preimplantation development, we analyzed published single-cell RNA-seq datasets from multiple eutherian mammals (human, rhesus monkey, marmoset, mouse, goat, cattle, pig) and the metatherian opossum. All RNA-seq reads were mapped to their corresponding genomes with retrotransposon expression summed at the subfamily level. Retrotransposon reads overlapping with annotated exons were excluded from our quantitation to avoid confounding gene and retrotransposon expression (Fig. 1A, 1B). For most mammalian species examined, retrotransposons collectively constitute one of the most abundant non-coding transcript species in preimplantation embryos, accounting for 7% to 34% of the transcriptome at peak expression across species (Fig. 1A, fig. S1A, table S1). Although retrotransposon sequences and integration sites are highly divergent among species, primate, livestock and mouse preimplantation embryos all exhibit a similar global retrotransposon expression profile, with a major switch at zygotic genome activation (ZGA) (Fig. 1A, 1B, fig. S1A, table S1). This global, dynamic retrotransposon expression profile closely resembles that of protein-coding genes, suggesting that retrotransposons are an integral component of mammalian preimplantation transcriptomes (Fig1B, fig. S1A, table S1 and S2).

**Fig. 1.**
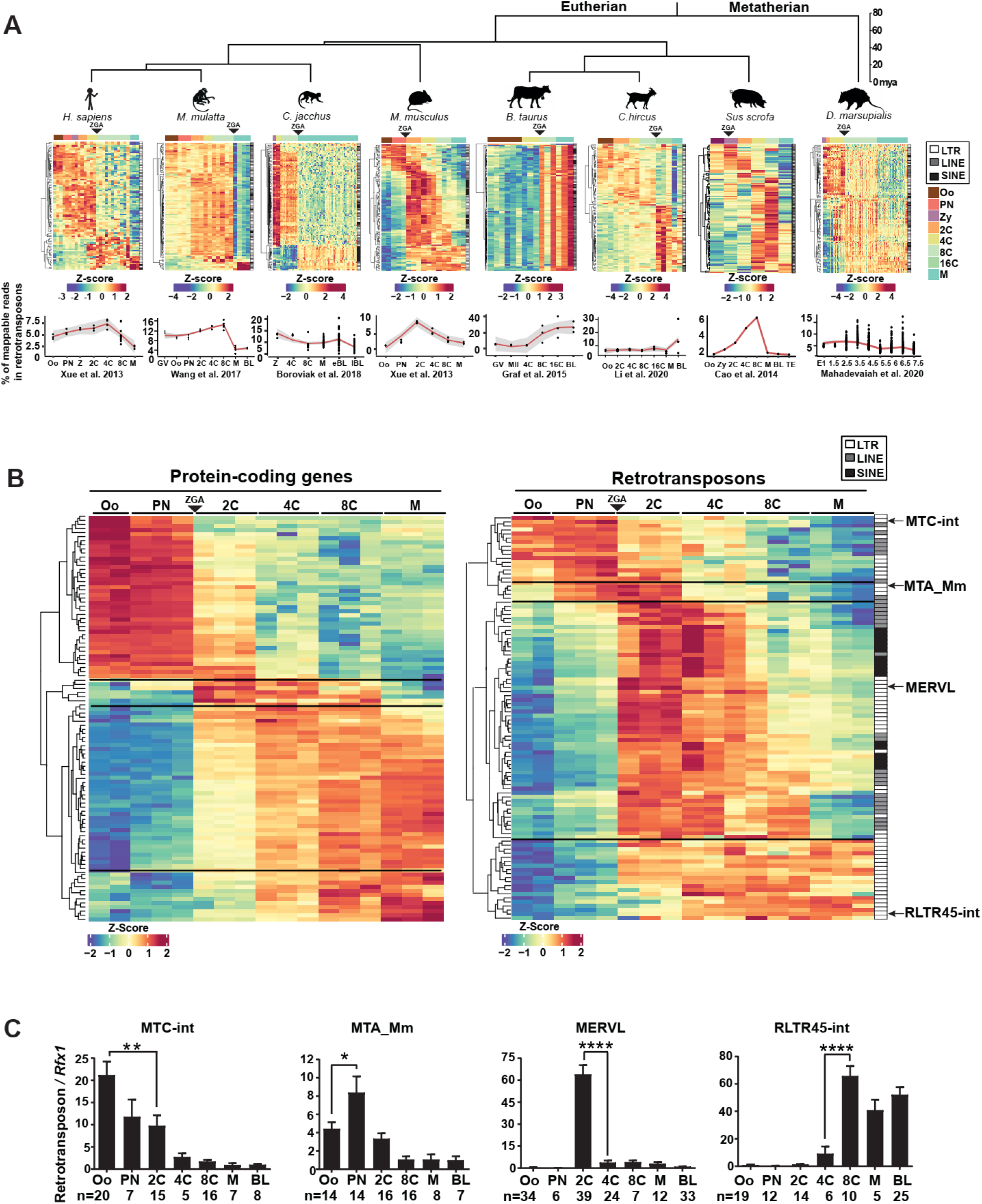
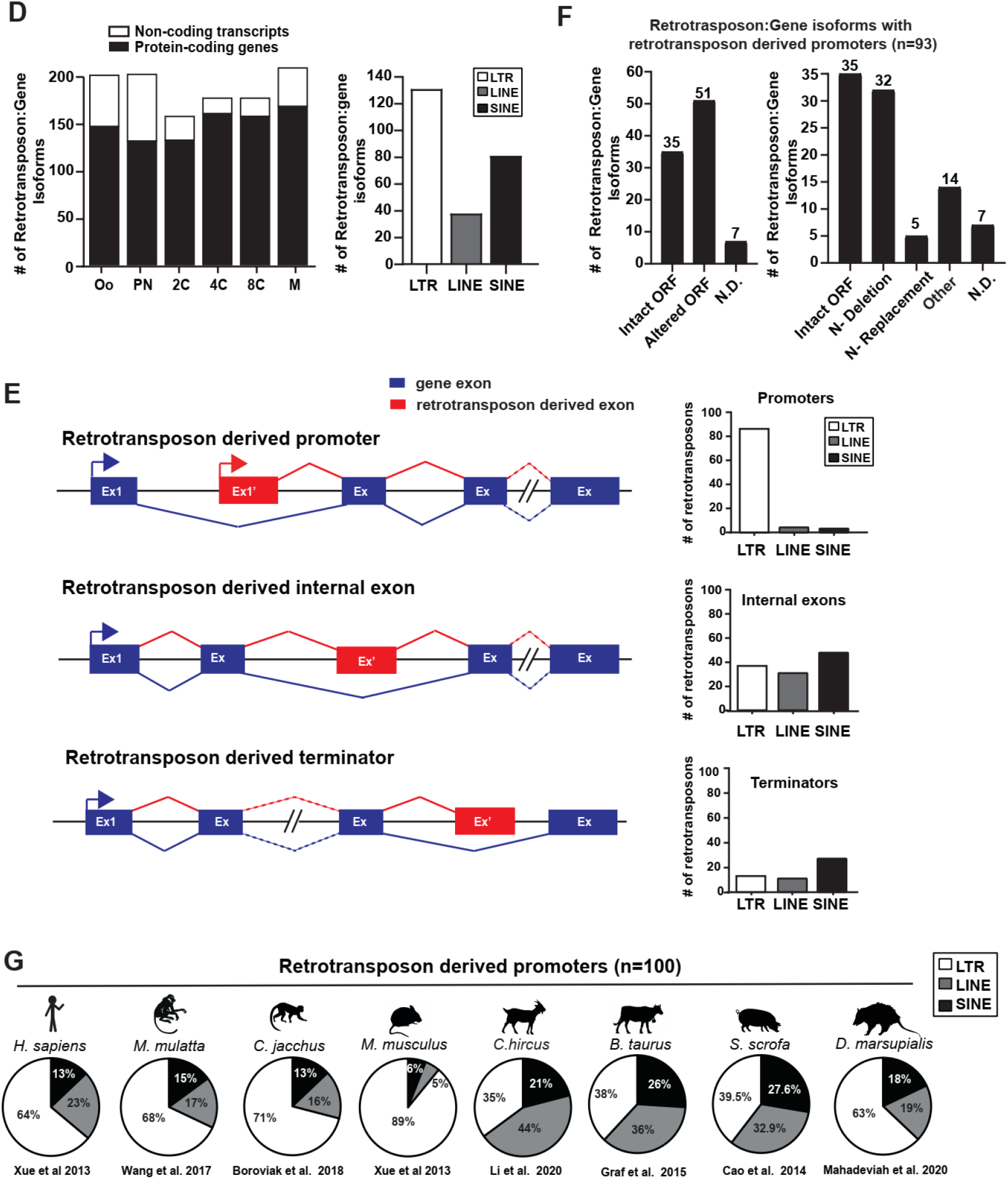
Retrotransposons mediate gene regulation in mammalian preimplantation development. **A**. Retrotransposons are highly and dynamically expressed in preimplantation embryos across multiple mammalian species, including human, rhesus monkey, marmoset, mouse, goat, cattle, pig and opossum. For each mammalian species, a heatmap exhibits the preimplantation profile of the top 100 most highly and differentially expressed retrotransposon subfamilies, and a line graph shows the percentage of uniquely mapped reads that originate from annotated retrotransposons. To highlight the comparison among species, only a subset of preimplantation stages was shown in heatmaps. All the developmental stages that are available in the original studies were included in line plots. **B**. Retrotransposons and protein-coding genes exhibit similar preimplantation expression profiles in mice, characterized by four distinct patterns. The top 100 most highly and differentially expressed protein-coding genes (left) and retrotransposon subfamilies (right) are shown as heatmaps. Representative subfamilies are marked with arrows. **A, B**. White, LTR; grey, LINE; black, SINE; black triangle, ZGA; Z-score, the number of standard deviations from the expression mean of a retrotransposon subfamily or a protein-coding gene. Oo, oocyte; Zy, zygotes; PN, pronucleus; 2C, two cell embryo; 4C, four cell embryo; 8C, eight cell embryo; 16C, sixteen cell embryos; M, morula; BL, blastocysts. **C**. Single embryo real time PCR analyses confirm the dynamic expression of four retrotransposon subfamilies, each representing a specific pattern. Error bars, ± s.e.m. *P* values were calculated using unpaired, two-tailed Student’s *t* test. (MTC-int, Oo vs. 2C, \*\**P* = 0.009, t=2.8, df=33 MTA_Mm, Oo vs. PN, \**P* = 0.04 t=2.1, df=26; MERVL, 2C vs. 4C, \*\*\*\**P* < 0.0001 t=7.4, df=62; RLTR45-int, 4C vs 8C, \*\*\*\**P* < 0.0001 t=5.2, df=16). **D**. Preimplantation-specific retrotransposon:gene splicing junctions preferentially associate with protein-coding genes across all preimplantation stages. Association of retrotransposon:gene splicing junctions with protein-coding genes vs. non-coding transcripts (left), and with LTR, LINE and SINE retrotransposon classes (right), are shown as bar plots. Only the most highly and differentially expressed retrotransposon:gene splicing junctions with ≥ 30 reads at each developmental stage are included in these analyses. **E**. Retrotransposons mediate gene regulation by acting as alternative promoters, internal exons and terminators for proximal gene isoforms. Representative gene structures are shown for retrotransposon-dependent gene isoforms (left). The top 250 most highly and differentially expressed retrotransposons that yield gene promoters (where TSS is within retrotransposon), internal exons and terminators were classified by LTRs, LINEs and SINEs (right). LTR retrotransposons are particularly enriched for promoters. **F**. Retrotransposon promoters frequently drive gene isoforms with N-terminally altered ORFs. Among the 250 most highly and differentially expressed retrotransposon:gene isoforms in mouse preimplantation embryos, 93 are driven by retrotransposon promoters. Manual curation of these 93 gene isoforms predicts frequent ORFs alterations (left), which are further classified based on the mechanisms of ORF alteration (right). N-Deletion, predicted N-terminal truncation; N-Replacement, predicted replacement of N-terminal protein sequence with retrotransposon derived sequences; Other, predicted with multiple N-terminal modification mechanisms; N.D., not determined. **G**. Retrotransposon derived gene promoters in preimplantation embryos are enriched for LTRs in mammalian genomes. The proportion of LTR, LINE and SINE retrotransposons was determined for the top 100 most highly and dynamically expressed retrotransposon promoters in preimplantation embryos of 8 mammalian species. In all species, LTR retrotransposons are significantly enriched. RNA-seq data for 1B, 1D, 1E and 1F analyses were obtained from Xue et al. 2013.

In mouse preimplantation embryos, retrotransposons exhibit four distinct expression patterns (Fig. 1B, 1C, fig. S1B and table S1), represented by MTC-int (induced in oocytes and silenced upon ZGA), MTA_Mm (a transient peak at pronuclear and 2C embryos), MERVL, ORR1A1 and IAPez-int (a transient peak in 2C-8C embryos), and RLTR45 and ERVB4_2-I_Mm (a peak in morula/blastocysts following the 8C induction) (Fig. 1C, table S1). Related retrotransposon families often exhibit similar expression patterns (data not shown), possibly due to shared ancestral transcriptional regulation(*46*–*48*).

Hundreds of preimplantation-specific splicing events are detected between a transcribed retrotransposon and a proximal gene exon (fig. S1C and table S3). Such retrotransposon-gene splicing events predominantly impact host protein-coding genes, rather than non-coding RNAs (Fig. 1D). Among the top 250 most highly and differentially expressed retrotransposon:gene isoforms, retrotransposons provide alternative promoters (37%), internal exons (46%) and terminators (17%) to proximal host genes (Fig. 1E, fig S1D). Using 5’RACE and real time PCR, we experimentally validated the gene structure and expression patterns of 27 predicted gene isoforms, with retrotransposons acting as alternative promoters, internal exons or terminators (table S4). Interestingly, a cohort of highly dynamic retrotransposon:gene isoforms differ from the corresponding canonical isoforms in gene structure, expression regulation, and in some cases, open reading frames (ORFs) (table S4, S5). Such alternative ORFs often harbor truncations, insertions or sequence replacement (fig. S1E), but rarely frame shift or non-sense mutations (fig. S1E, table S5). Hence, a selective pressure likely acts to preserve protein coding retrotransposon:gene isoforms with specific functions.

Retrotransposon-derived promoters have emerged as important players in development and disease(*7, 38, 49*–*51*), often driving retrotransposon:gene isoforms. Retrotransposon promoters in mouse preimplantation embryos are particularly enriched for LTR retrotransposons, rather than LINEs or SINEs (Fig. 1E). Sequence related LTR promoters, when simultaneously induced, appear to coordinately transcribe a cohort of host genes (fig. S1F). LTR retrotransposons exist either as full proviral sequences with two identical long terminal repeats (LTRs) flanking an internal region, or more frequently, as solo-LTRs (fig. S1G). The LTR retrotransposons could retain viral promoter sequences, not only conferring new transcriptional regulation, but also frequently contributing to alternative 5’UTRs and/or modifying ORFs of proximal host genes (Fig. 1F). We manually curated the top 93 highly and differentially expressed retrotransposon promoters in mouse preimplantation embryos and revealed that 55% were predicted to drive alternative gene isoforms that encode N-terminally altered ORFs (Fig. 1F, table S5).

In addition to mouse, human, rhesus monkey, macaque, cattle, goat, pig and opossum all employ numerous retrotransposon promoters in preimplantation embryos. Despite their sequence divergence in different species, the LTR class is enriched in retrotransposon promoters across all mammalian preimplantation embryos examined (Fig. 1G, fig. S1H, tables S3 and S5).

The prevalence of retrotransposon initiated preimplantation gene isoforms prompted us to explore its functional importance *in vivo*. One of the most highly and dynamically expressed, mouse retrotransposon:gene isoform is driven by an MT2B2 LTR retrotransposon promoter (table S3). The MT2B2 promoter, located 8.2 kb upstream of *Cdk2ap1* (Cyclin dependent kinase associated protein), drives an N-terminally truncated *Cdk2ap1* isoform, *Cdk2ap1*^*ΔN(MT2B2)*^ (Fig. 2A, tables S3, S4 and S5).

**Fig. 2.**
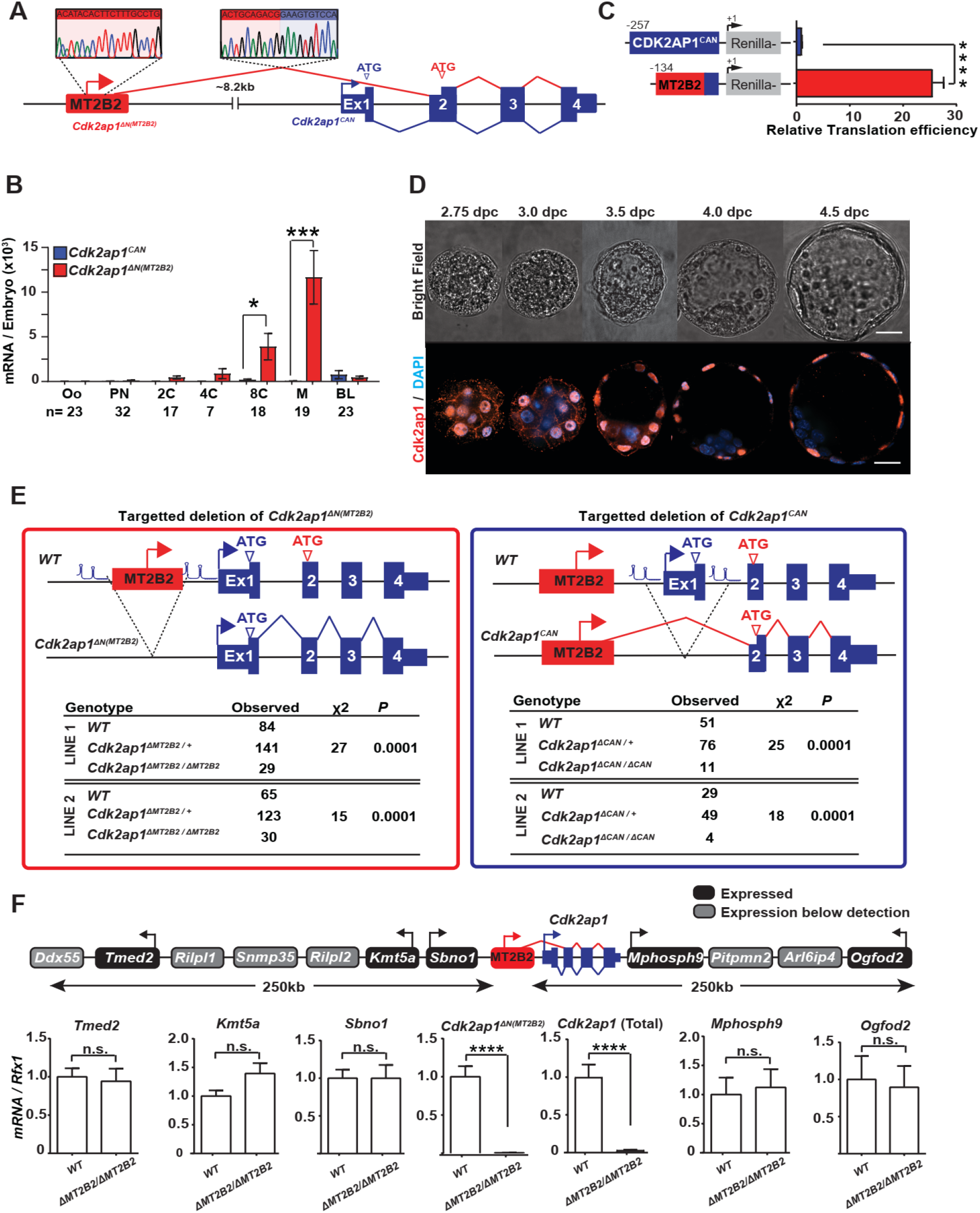
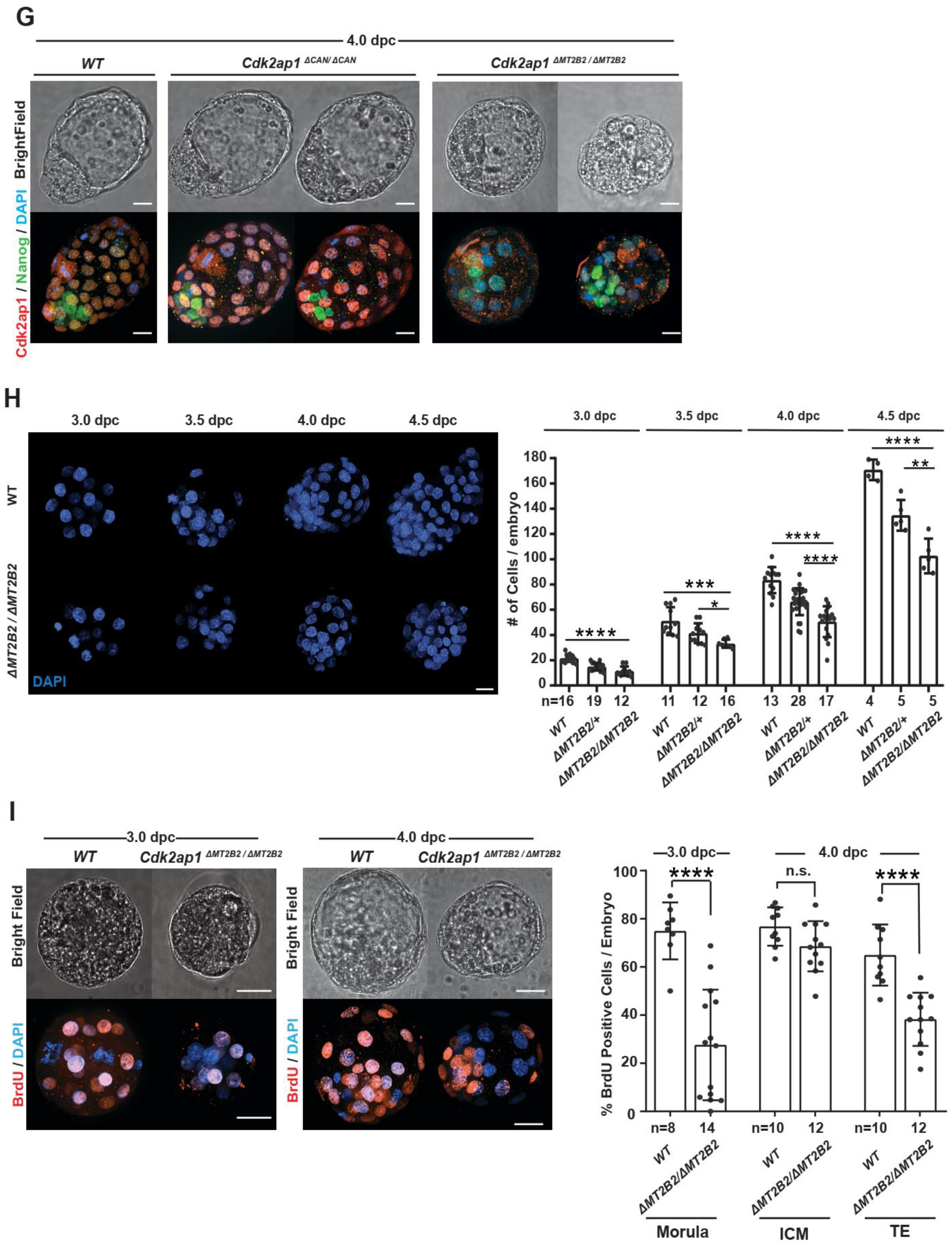
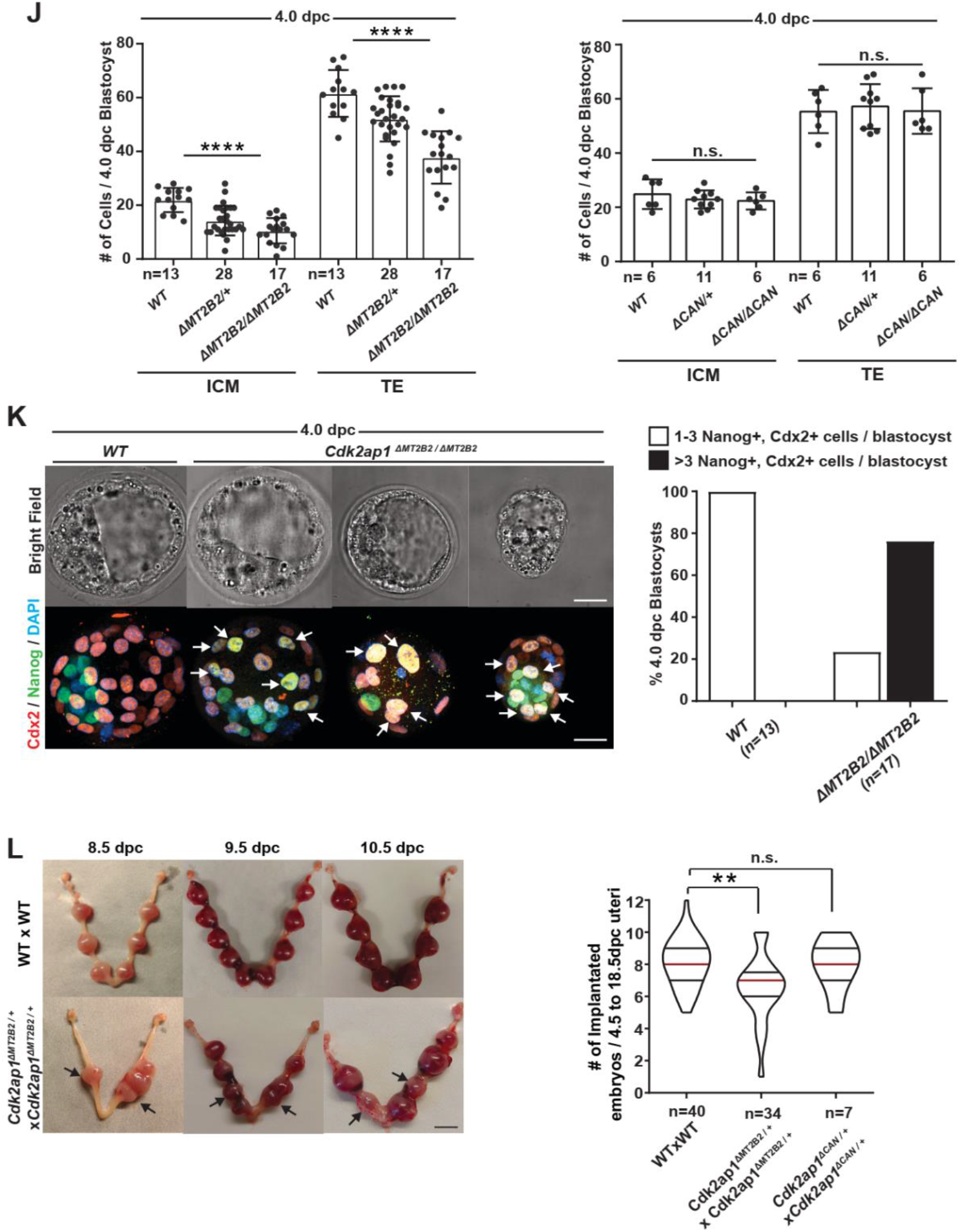
An MT2B2 retrotransposon promoter drives an N-terminally truncated *Cdk2ap1*^*ΔN(MT2B2)*^ isoform that is essential for preimplantation development. **A**. A diagram illustrates the gene structure of the canonical *Cdk2ap1*^*CAN*^ isoform (blue) and the preimplantation-specific *Cdk2ap1*^*ΔN(MT2B2)*^ isoform (red). 5’ RACE confirms the transcription start site within the MT2B2 element; RT-PCR and Sanger sequencing confirm the splicing between the MT2B2 element and the *Cdk2ap1* exon 2. **B**. Absolute real-time PCR quantification of single embryos compare the level of *Cdk2ap1*^*CAN*^ and *Cdk2ap1*^*ΔN(MT2B2)*^ isoforms in mouse preimplantation embryos. Error bars, s.e.m. *Cdk2ap1*^*CAN*^ vs. *Cdk2ap1*^*ΔN(MT2B2)*^ at 8C stage, n=17, \**P* = 0.02, t=2.5, df=34; *Cdk2ap1*^*CAN*^ vs. *Cdk2ap1*^*ΔN(MT2B2)*^ at morula stage, n=19, \*\*\**P* = 0.0004, t=3.9, df=36. **C**. The MT2B2 derived 5’UTR enhances the translation efficiency of the *Cdk2ap1*^*ΔN(MT2B2)*^ isoform. The 5’UTRs of *Cdk2ap1*^*CAN*^ and *Cdk2ap1*^*ΔN(MT2B2)*^ were each cloned 5’ to a Renilla luciferase reporter, transfected in HEK293T cells, and assayed for translation efficiency using luciferase assays. The MT2B2 derived 5’UTR sequence in *Cdk2ap1*^*ΔN(MT2B2)*^ was associated with a higher translation efficiency. Three independent experiments were performed, and transfections were performed in triplicates per condition. Error bars, s.e.m; **** *P* < 0.0001, t=20.44, df=4. **D**. Mouse preimplantation embryos between 2.75 dpc to 4.5 dpc were subjected to immunostaining for Cdk2ap1. Cdk2ap1 protein exhibits specific expression in the outer cells of morulae and the TE cells in blastocysts. Confocal images are representative of 4 or more embryos per stage. Scale bar, 20µm. **E**. Diagrams illustrate the CRISPR genome engineering strategy for specific targeted deletion of *Cdk2ap1*^*ΔN(MT2B2)*^ (left) and *Cdk2ap1* (right) in mice. Mendelian ratio of progenies from the *Cdk2ap1*^*ΔMT2B2/+*^ x *Cdk2ap1*^*ΔMT2B2/+*^ crosses (left) and the *Cdk2ap1*^*ΔCAN/+*^ x *Cdk2ap1*^*ΔCAN/+*^ (right) was documented at postnatal day 10 (p10), demonstrating a significant reduction of viability in both genotypes. Two independent *Cdk2ap1*^*ΔMT2B2/ΔMT2B2*^ lines and two independent *Cdk2ap1*^*ΔCAN/ΔCAN*^ lines were subjected to these analyses. **F**. The MT2B2 deletion specifically abolishes *Cdk2ap1*^*ΔN(MT2B2)*^ expression, but not neighboring genes. Age matched wildtype (n=9) and *Cdk2ap1*^*ΔMT2B2/ΔMT2B2*^ (n=7) morula embryos were collected from two independent WT x WT and *Cdk2ap1*^*ΔMT2B2/ΔMT2B2*^ x *Cdk2ap1*^*ΔMT2B2/ΔMT2B2*^ crosses, and subjected to single embryo real-time PCR analyses to measure the expression of *Cdk2ap1*^*ΔN(MT2B2)*^, total *Cdk2ap1* and all neighboring genes with 250 kb of the deletion. Black, expressed genes; grey, genes below detection; error bars, s.e.m. *Cdk2ap1*(Total), wildtype (n=3) vs. *Cdk2ap1*^*ΔMT2B2/ΔMT2B2*^ (n=3), \*\*\*\**P* < 0.0001, t=16.8, df=4; *Cdk2ap1*^*ΔN(MT2B2)*^, wildtype (n=9) vs. *Cdk2ap1*^*ΔMT2B2/ΔMT2B2*^ (n=7), \*\*\**P* = 0.0002, t=4.9, df=14. **G**. *Cdk2ap1*^*ΔMT2B2/ΔMT2B2*^ embryos, but not *Cdk2ap1*^*ΔCAN/ΔCAN*^ embryos, exhibit defective Cdk2ap1 protein expression in TE and impaired blastocyst formation. Representative confocal images shown for wildtype (n=11), *Cdk2ap1*^*ΔCAN/ΔCAN*^ (n=5) and *Cdk2ap1*^*ΔMT2B2/ΔMT2B2*^ (n=6) embryos subjected to immunostaining of Cdk2ap1 and Nanog. Scale bar, 25 μm. **H**. *Cdk2ap1*^*ΔMT2B2/ΔMT2B2*^ embryos exhibited reduced cell number during preimplantation development. A total of 158 embryos from littermate controlled wildtype (n=44), *Cdk2ap1*^*ΔMT2B2/+*^ (n= 64) and *Cdk2ap1*^*ΔMT2B2/ΔMT2B2*^ (n=50) embryos were collected at 3.0 dpc, 3.5 dpc, 4.0 dpc and 4.5 dpc from a total of 29 *Cdk2ap1*^*ΔMT2B2/+*^ to *Cdk2ap1*^*ΔMT2B2/+*^ matings. Representative images of DAPI staining (left) and cell number quantitation (right) are shown for each developmental stage. Scale bar, 25 μm; error bars, s.d.. Wildtype vs. *Cdk2ap1*^*ΔMT2B2/ΔMT2B2*^ embryos: 3.0 dpc, **** *P* < 0.0001, t=8.2, df=26; 3.5 dpc, *** *P* = 0.0007, t=4.2, df=15; 4.0 dpc, **** *P* < 0.0001, t=7.8, df=28; 4.5 dpc, **** *P* < 0.0001, t=8.7, df=7. *Cdk2ap1*^*ΔMT2B2/+*^*vs Cdk2ap1*^*ΔMT2B2/ΔMT2B*^, 3.5 dpc, * *P* = 0.03, t=2.4, df=16; 4.0 dpc, **** *P* < 0.0001, t=4.5, df=43; 4.5 dpc, ** *P* = 0.005, t=3.9 df=8. **I**. *Cdk2ap1*^*ΔMT2B2/ΔMT2B2*^ embryos exhibit decreased S-phase entry in BrdU incorporation assays. Representative confocal images (left) and quantitation (right) of BrdU staining are shown for embryos at 3.0 and 4.0 dpc. Age matched wildtype (n=18) and *Cdk2ap1*^*ΔMT2B2/ΔMT2B2*^ (n=26) morulae and blastocysts were collected from three wildtype x wildtype and six *Cdk2ap1*^*ΔMT2B2/ΔMT2B2*^ x *Cdk2ap1*^*ΔMT2B2/ΔMT2B2*^ matings, respectively. Scale bars, 20 μm; error bars, s.d.. Wildtype vs. *Cdk2ap1*^*ΔMT2B2/ΔMT2B2*^ embryos: morulae, **** *P* < 0.0001, t=7.9, df=20; TE, **** *P* < 0.0001, t=5.3, df=20. **J**. *Cdk2ap1*^*ΔMT2B2/ΔMT2B2*^ blastocysts, but not *Cdk2ap1*^*ΔCAN/ΔCAN*^ blastocysts, exhibit a decreased cell number in ICM and TE. Littermate controlled wildtype and knockout blastocysts were subjected to immunostaining of Nanog and Cdx2 for the quantitation of ICM cells and TE cells, respectively. Scale bars, 25 μm; error bars, s.d.. Wildtype vs *Cdk2ap1*^*ΔMT2B2/ΔMT2B2*^: ICM, **** *P* < 0.0001, t=6.7, df=28; TE, **** *P* < 0.0001, t=6.9, df=28. **K**. The MT2B2 deletion impairs cell fate specification in blastocysts. Littermate controlled wildtype (n=13) and *Cdk2ap1*^*ΔMT2B2/ΔMT2B2*^ (n=17) blastocysts were collected at 4.0 dpc and subjected to immunostaining of Nanog and Cdx2. The presence of ≥ 3 Nanog and Cdx2 double positive cells in *Cdk2ap1*^*ΔMT2B2/ΔMT2B2*^ suggests an impaired cell fate specification. Representative confocal images (left) and quantitation (right) were shown for Nanog and Cdx2 double positive cells in each genotype. White arrows, Nanog and Cdx2 double positive cells. Scale bar 0.5 cm. **L**. The deletion of *Cdk2ap1*^*ΔN(MT2B2)*^, but not *Cdk2ap1*^*CAN*^, causes aberrant embryo spacing and defective embryo implantation. Representative images were shown for embryo implantation at 8.5, 9.5 and 10.5 dpc in wildtype x wildtype and *Cdk2ap1*^*ΔMT2B2/+*^ x *Cdk2ap1*^*ΔMT2B2/+*^ crosses (left). Black arrows, genotype confirmed *Cdk2ap1*^*ΔMT2B2/ΔMT2B2*^ embryos; scale bar, 0.5 cm. Quantitation of implanted embryos from 4.5 to 18.5 dpc per uterus was shown for wildtype x wildtype (n=40 uteri), *Cdk2ap1*^*ΔMT2B2/+*^ x *Cdk2ap1*^*ΔMT2B2/+*^ (n=34 uteri) and *Cdk2ap1*^*ΔCAN/+*^ x *Cdk2ap1*^*ΔCAN/+*^ (n=7 uteri) crosses as a violin plot, with median (red line) as well as lower (25%) and upper (75%) quartiles (black lines). Wildtype x wildtype vs. *Cdk2ap1*^*ΔMT2B2/+*^ x *Cdk2ap1*^*ΔMT2B2/+*^, ** *P* = 0.002, t=3.2, df=72. All *P* values were calculated using unpaired, two-tailed Student’s *t* test, unless otherwise stated. n.s., not significant.

The canonical *Cdk2ap1* (*Cdk2ap1*^*CAN*^) is characterized as a suppressor of cell proliferation, at least in part, by promoting Cdk2 degradation and repressing kinase activity(*52*–*54*). For *Cdk2ap1*^*CAN*^, both transcription start site (TSS) and the ATG start codon are found within exon 1 (Fig. 2A). In contrast, the MT2B2 driven *Cdk2ap1*^*ΔN(MT2B2)*^ isoform is alternatively spliced to skip exon 1 and utilizes a downstream ATG found in exon 2, resulting in an N-terminal truncation of the first 27 amino acids (Fig. 2A, fig. S2A). The MT2B2 element not only promotes strong *Cdk2ap1*^*ΔN(MT2B2)*^ induction at the 8C to morula stages (Fig. 2B), but also contributes to a hybrid 5’UTR sequence with enhanced translation efficiency (Fig. 2C).

*Cdk2ap1*^*CAN*^ and *Cdk2ap1*^*ΔN(MT2B2)*^ exhibit distinct expression patterns in mouse embryonic development. *Cdk2ap1*^*CAN*^ remained at a low level throughout preimplantation development, and later peaked around 10.5 days post coitum (10.5 dpc) (Fig. 2B and fig. S2B). In comparison, *Cdk2ap1*^*ΔN(MT2B2)*^ is the predominant preimplantation-specific *Cdk2ap1* transcript, with peak expression from 8C to morula embryos (Fig. 2B). Cdk2ap1 protein expression was slightly delayed, first detected in the nuclei of compacted morula blastomeres, and subsequently in the trophectoderm (TE) cells of blastocyst (Fig. 2D). This is consistent with its mRNA enrichment in the TE cells of blastocyst embryos (fig. S2C).

To determine which Cdk2ap1 protein isoform is expressed in preimplantation embryos, we engineered isoform-specific endogenous V5 tagging at the N-terminus of *Cdk2ap1*^*CAN*^, the N-terminus of *Cdk2ap1*^*ΔN(MT2B2)*^, and the C-terminus of all *Cdk2ap1* isoforms. V5 Immunostaining revealed that most, if not all, Cdk2ap1 protein in preimplantation embryos is generated from *Cdk2ap1*^*ΔN(MT2B2)*^ (fig. S2D).

We next investigated the functional importance of the MT2B2 retrotransposon promoter. We employed CRISPR-EZ, a highly efficient CRISPR technology for mouse genome engineering(*55, 56*). We deleted the MT2B2 element or the *Cdk2ap1* canonical exon 1, generating C57B/6J mice deficient for either *Cdk2ap1*^*ΔN(MT2B2)*^ or *Cdk2ap1*^*CAN*^, respectively (designated as *Cdk2ap1*^*ΔMT2B2/ΔMT2B2*^ and *Cdk2ap1*^*ΔCAN/ΔCAN*^ mice, Fig. 2E, fig. S2E). The MT2B2 deletion specifically abolished *Cdk2ap1*^*ΔN(MT2B2)*^, and significantly reduced total *Cdk2ap1* mRNA and protein in preimplantation embryos without impacting any flanking genes within 250kb (Fig. 2F, 2G). While both *Cdk2ap1*^*ΔMT2B2/ΔMT2B2*^ and *Cdk2ap1*^*ΔCAN/ΔCAN*^ mice exhibited significantly reduced viability at P10 (Fig. 2E), only *Cdk2ap1*^*ΔMT2B2/ΔMT2B2*^ mice exhibited preimplantation defects (Fig. 2G).

Two independent *Cdk2ap1*^*ΔMT2B2/ΔMT2B2*^ mouse lines exhibited 50-55% penetrance for lethality by postnatal day 10 (p10) (Fig. 2E); those that survive into adulthood appear grossly normal and are fertile (data not shown). *Cdk2ap1*^*ΔMT2B2/ΔMT2B2*^ embryos were recovered at the expected Mendelian ratio at 4.0 dpc, yet 71% (12/17) exhibited abnormal morphology, characterized by reduced cell number, aberrant cell organization and impaired blastocoel cavities (Fig. 2G, 2H). In particular, *Cdk2ap1*^*ΔN(MT2B2)*^ deficiency significantly reduced cell proliferation in preimplantation embryos, as a decrease in both total cell number and BrdU incorporation was observed in morula and blastocyst embryos (Fig. 2H, 2I and 2J), particularly affecting the TE compartment. This finding suggested an impaired S-phase entry caused by the MT2B2 deletion (Fig. 2I). Consistently, aberrant Nanog and Cdx2 double-positive cells were frequently identified in *Cdk2ap1*^*ΔMT2B2/ΔMT2B2*^ blastocysts, indicating a delayed/impaired cell fate specification (Fig. 2K). Reduced TE cell number and impaired TE cell fate specification in *Cdk2ap1*^*ΔMT2B2/ΔMT2B2*^ blastocysts likely contribute to decreased implantation rate, aberrant embryo spacing in uterus, as well as increased embryo lethality in development (Fig. 2L and fig. S2F). The blastocyst defects observed in *Cdk2ap1*^*ΔMT2B2/ΔMT2B2*^ embryos bear phenotypic resemblance to the preimplantation lethality caused by targeted disruption of all *Cdk2ap1* isoforms (*57*).

To our knowledge, MT2B2 is the first example of a retrotransposon promoter with an essential function in mammalian development. Previous studies have characterized the knockout phenotype of two retrotransposon promoters in mice and flies, yet the resulted defects affect mating behavior and female fertility, both of which are non-essential for normal development (*25, 58*).

In contrast to *Cdk2ap1*^*ΔMT2B2/ΔMT2B2*^ embryos, *Cdk2ap1*^*ΔCAN/ΔCAN*^ blastocysts were morphologically intact, with no defects in cell number, cell proliferation, or cell fate specification (Fig. 2G, 2J). Nevertheless, two independent *Cdk2ap1*^*ΔCAN/ΔCAN*^ mouse lines both exhibited reduced viability by p10, with a 58-67% penetrance for lethality (Fig. 2E). Implanted *Cdk2ap1*^*ΔCAN/ΔCAN*^ embryos exhibited an increase in resorption events (fig. S2F), in contrast to *Cdk2ap1*^*ΔMT2B2/ΔMT2B2*^ embryos that instead frequently show implantation spacing defects (Fig. 2L). The lethality of *Cdk2ap1*^*ΔCAN/ΔCAN*^ mice is likely attributed to defects in post-implantation development, consistent with the peak expression of *Cdk2ap1*^*CAN*^ during mid-gestation (fig. S2B).

The effect of *Cdk2ap1*^*ΔN(MT2B2)*^ on cell proliferation is opposite from the anti-proliferative function described for *Cdk2ap1*^*CAN*^ (*52, 59, 60*). The decreased blastomere count in *Cdk2ap1*^*ΔMT2B2/ΔMT2B2*^ embryos supports a role for *Cdk2ap1*^*ΔN(MT2B2)*^ in promoting cell proliferation. To investigate their opposite effects on cell proliferation, we compared *Cdk2ap1*^*ΔN(MT2B2)*^ and *Cdk2ap1*^*CAN*^ in their overexpression phenotypes in wildtype and *Cdk2ap1*^*ΔMT2B2/ΔMT2B2*^ embryos. We optimized an electroporation-based method for mRNA delivery into mouse zygotes (Fig. 3A, fig. S3A), and achieved an mRNA delivery efficiency comparable to that by mRNA microinjection (data not shown). Wildtype zygotes overexpressing *Cdk2ap1*^*ΔN(MT2B2)*^ developed into embryos with both increased cell number and greater BrdU incorporation (Fig. 3B and fig. S3B). Conversely, those overexpressing *Cdk2ap1*^*CAN*^ gave rise to embryos with reduced BrdU incorporation and decreased cell numbers (Fig. 3B and fig. S3B). Importantly, ectopic *Cdk2ap1*^*ΔN(MT2B2)*^ expression rescued *Cdk2ap1*^*ΔMT2B2/ΔMT2B2*^ embryos, restoring BrdU incorporation and total cell number to wildtype levels, and mitigating Nanog and Cdx2 double positivity in blastocysts (Fig. 3C, 3D, 3E and fig. S3B). In contrast, *Cdk2ap1*^*CAN*^ overexpression in *Cdk2ap1*^*ΔMT2B2/ΔMT2B2*^ embryos exacerbated cell proliferation and cell fate defects (Fig. 3C, 3D, 3E and fig. S3B). Hence, *Cdk2ap1*^*ΔN (MT2B2)*^ and *Cdk2ap1*^*CAN*^ exhibit opposite effects on cell proliferation in preimplantation embryos, but functional antagonism is unlikely due to non-overlapping expression patterns.

**Fig. 3.**
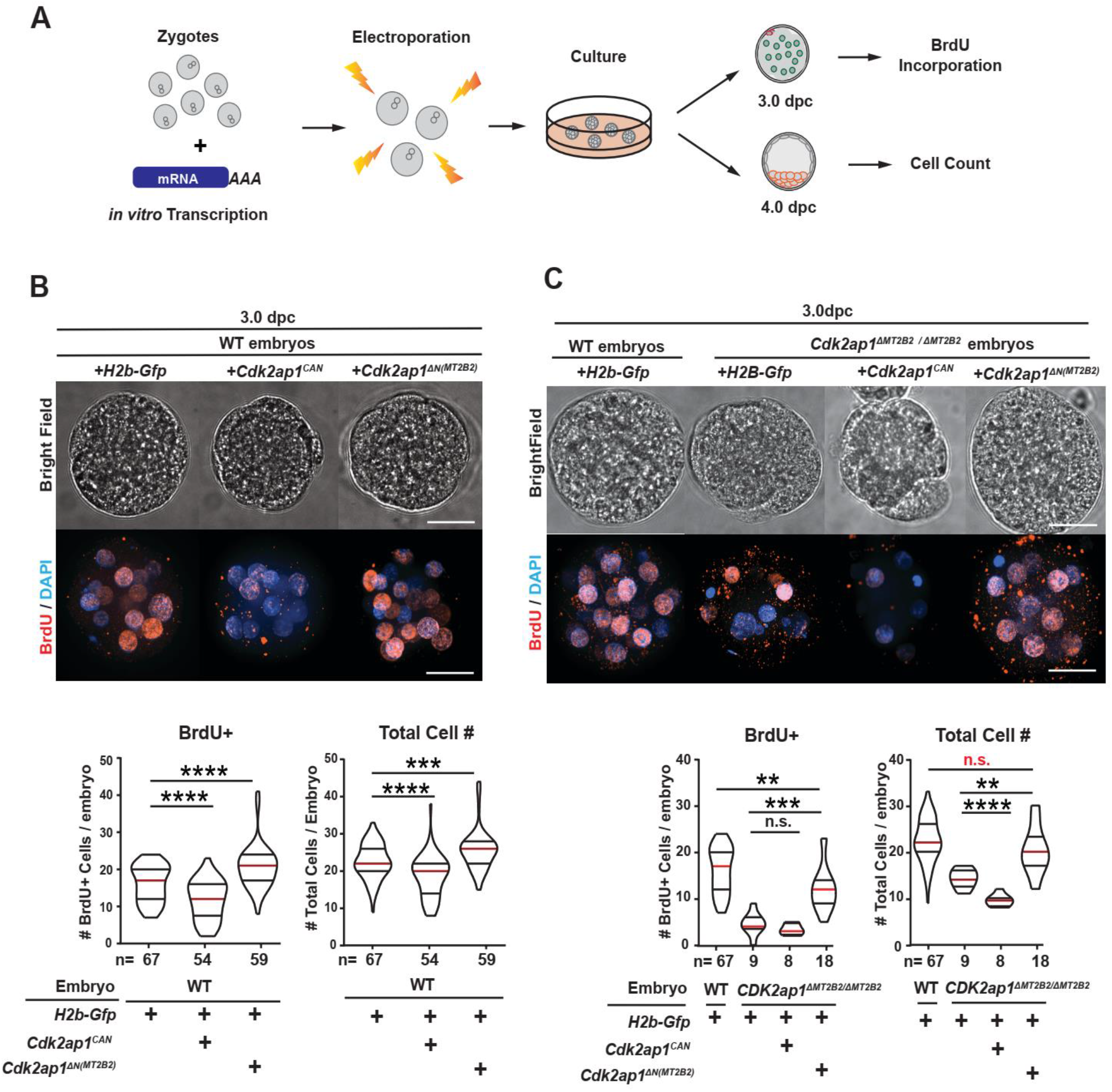
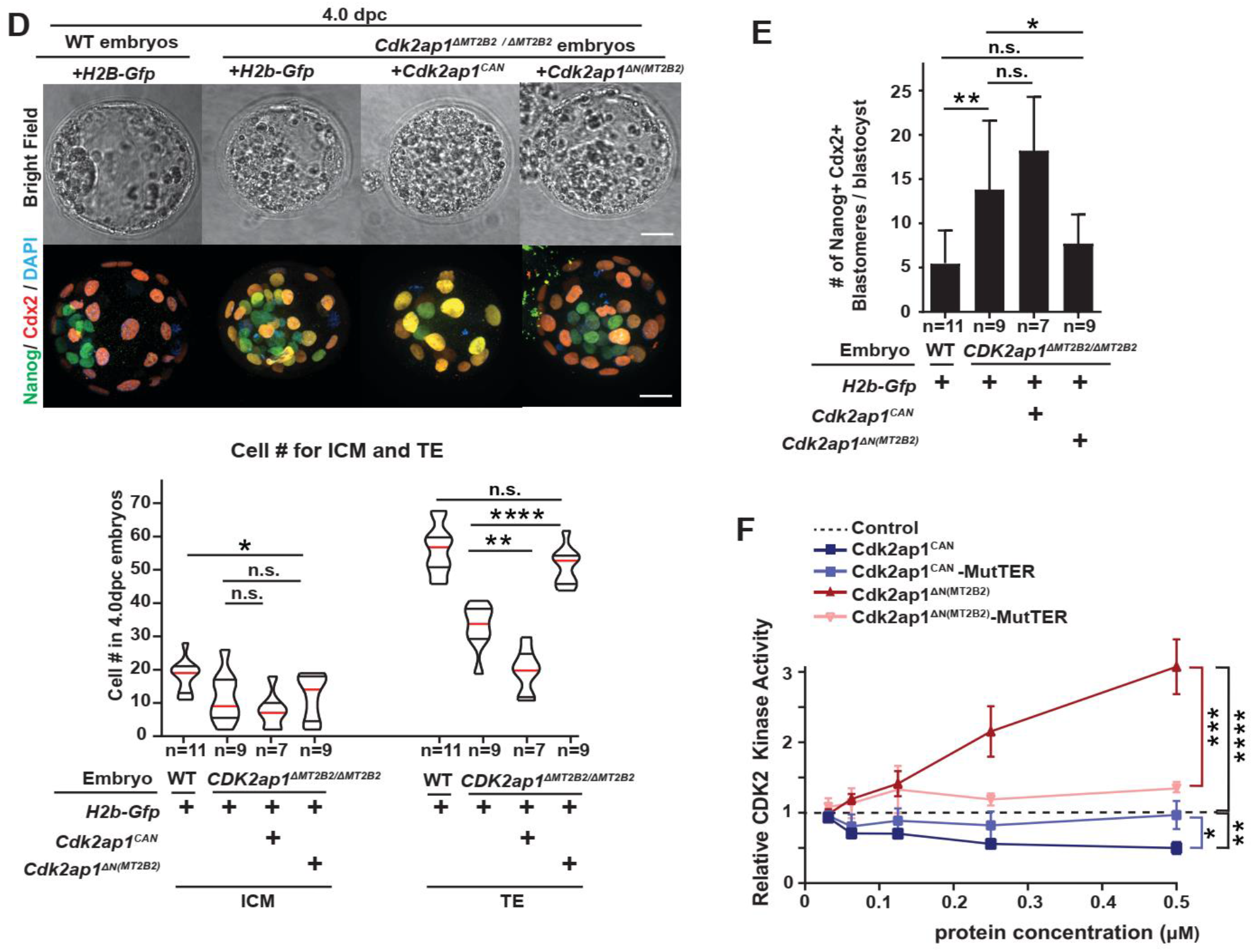
Cdk2ap1^ΔN(MT2B2)^ and Cdk2ap1^CAN^ have opposite functions in cell proliferation. **A**. A diagram illustrates the experimental scheme for mRNA delivery into zygotes using electroporation. **B, C**. The *Cdk2ap1*^*CAN*^ and the *Cdk2ap1*^*ΔN(MT2B2)*^ isoforms have opposite effects on S-Phase entry and cell proliferation in wildtype and *Cdk2ap1*^*ΔMT2B2/ΔMT2B2*^ embryos. *H2b-Gfp, Cdk2ap1*^*CAN*^ or *Cdk2ap1*^*ΔN(MT2B2)*^ mRNAs were each electroporated into wildtype (**B**) or *Cdk2ap1*^*ΔMT2B2/ΔMT2B2*^ (**C**) pronuclear embryos, and resulted embryos were compared for BrdU incorporation in morulae at 3.0 dpc. Ectopic expression of *Cdk2ap1*^*ΔN(MT2B2)*^ restores S-Phase entry and cell proliferation in *Cdk2ap1*^*ΔMT2B2/ΔMT2B2*^ embryos at 3.0 dpc. Representative immunostaining images (top) and quantitation of BrdU positive and total cell number (bottom) were shown. Violin plots were shown with median (red), as well as lower (25%) and upper (75%) quartiles (black lines). Scale bars, 20 μm. *H2b-Gfp* vs *Cdk2ap1*^*CAN*^ in wildtype embryos: BrdU, **** *P* < 0.0001, t=4.9, df=119; total cell number, **** *P* < 0.0001, t=4.1, df=119. *H2b-Gfp* vs *Cdk2ap1*^*ΔN(MT2B2)*^ in wildtype embryos: BrdU+, **** *P* < 0.0001, t=4.7, df=124; total cell number, *** *P* = 0.0005, t=3.6, df=124. *H2b-Gfp* vs *Cdk2ap1*^*CAN*^ in *Cdk2ap1*^*ΔMT2B2/ΔMT2B2*^ embryos: BrdU, n.s.; total cell number, **** *P* < 0.0001, t=5.7, df=15. *H2b-Gfp* vs *Cdk2ap1*^*ΔN(MT2B2)*^ in *Cdk2ap1*^*ΔMT2B2/ΔMT2B2*^ embryos: BrdU, *** *P* =0.0002, t=4.5, df=25; total cell number, ** *P* =0.002, t=3.4, df=25. **D, E**. Ectopic expression of *Cdk2ap1*^*ΔN(MT2B2)*^ rescues cell proliferation and cell fate specification defects in *Cdk2ap1*^*ΔMT2B2/ΔMT2B2*^ embryos. **D**. Representative confocal image of Cdx2 and Nanog immunostaining (top) and quantitation of ICM and TE cell number (bottom) were shown for 4.0 dpc *Cdk2ap1*^*ΔMT2B2/ΔMT2B2*^ embryos overexpressing *Cdk2ap1*^*CAN*^ or *Cdk2ap1*^*ΔN(MT2B2)*^. Scale bars 20 μm; White arrows, Nanog and Cdx2 double positive cells. *H2b-Gfp* vs *Cdk2ap1*^*CAN*^, TE, ** *P* =0.002, t=3.9, df=14; *H2b-Gfp* vs *Cdk2ap1*^*ΔN(MT2B2)*^, TE, **** *P* < 0.0001, t=6.1, df=16. **E**. Quantitation of Nanog and Cdx2 double positive cells was shown for *Cdk2ap1*^*ΔMT2B2/ΔMT2B2*^ embryos overexpressing *Cdk2ap1*^*CAN*^ or *Cdk2ap1*^*ΔN(MT2B2)*^. *H2b-Gfp*-overexpressing wildtype vs. *Cdk2ap1*^*ΔMT2B2/ΔMT2B2*^ embryos, ** *P* = 0.007, t=3.1, df=17; *H2b-Gfp* vs *Cdk2ap1*^*ΔN(MT2B2)*^ in *Cdk2ap1*^*ΔMT2B2/ΔMT2B2)*^ embryos, * *P* =0.04, t=2.2, df=16. **F**. Cdk2ap1^CAN^ and *Cdk2ap1*^*ΔN(MT2B2)*^ have opposite effects on Cdk2 kinase activity. We constructed Cdk2ap1 mutants (Cdk2ap1^CAN^- MutTER and Cdk2ap1^ΔN(MT2B2)^-MutTER) that fail to bind to Cdk2 by mutating the well-defined TER motif, the Cdk2 binding motif (T108A, E109A, R110A). Recombinant Cdk2ap1^CAN^- MutTER and Cdk2ap1^ΔN(MT2B2)^-MutTER proteins were purified by size separation column purification, before incubated with recombinant CDK2, CYCLIN E, HISTONE H1 and ATP *in vitro*. Their effects on CDK2 activity at different concentrations were analyzed in a kinase assay. Three independent experiments were performed. Dashed line, baseline CDK2 kinase activity when elution buffer is used as “control” input. Error bars, s.e.m. Control vs Cdk2ap*1*^*CAN*^, \*\**P* = 0.001, t=8.4, df=4. Cdk2ap*1*^*CAN*^ vs Cdk2ap*1*^*CAN*^-MutTER, \**P* = 0.02, t=3.8, df=4. Control vs *Cdk2ap1*^*ΔN(MT2B2)*^, \*\*\*\**P* < 0.0001, t=10.5, df=6; *Cdk2ap1*^*ΔN(MT2B2)*^ vs *Cdk2ap1*^*ΔN(MT2B2)*^*-*MutTER, \*\*\**P* = 0.0003, t=8.9, df=5. All *P* values were calculated using unpaired, two-tailed Student’s *t* test, unless otherwise stated. n.s., not significant.

Previous studies described Cdk2ap1^CAN^ as a potent, negative cell cycle regulator that directly binds to Cdk2 via a TER motif that reduces its abundance and inhibits kinase activity(*52, 59, 60*). Both Cdk2ap1^CAN^ and Cdk2ap1^*Δ*N(MT2B2)^ contain the TER motif and directly associate with Cdk2 in co-immunoprecipitation experiments (fig. S3C). The effects of Cdk2ap1^CAN^ and Cdk2ap1^*Δ*N(MT2B2)^ on Cdk2 activity were compared using an *in vitro* CDK2 Kinase assay. Immuno-precipitated Cdk2ap1 lysate from *Cdk2ap1*^*CAN*^ or *Cdk2ap1*^*ΔN(MT2B2)*^ overexpressing HEK293T cells were incubated with recombinant CDK2/CYCLIN E1 complex to quantify their effects on CDK2 kinase activity (fig S3D). Similarly, purified recombinant Cdk2ap1^CAN^ and Cdk2ap1^*Δ*N(MT2B2)^ proteins were tested in the same assay for their effects on CDK2 kinase activity (Fig. 3E). In both experiments, Cdk2ap1^CAN^ and Cdk2ap1^*Δ*N (MT2B2)^ significantly inhibited and enhanced CDK2 kinase activity, respectively (Fig. 3E, fig. S3D), in line with their opposite effects on cell proliferation *in vivo*. Mutation of the TER motif in Cdk2ap1^CAN^ or Cdk2ap1^*Δ*N(MT2B2)^ abolished their effects on CDK2 kinase activity (Fig. 3F, fig. S3D), demonstrating the importance of direct Cdk2ap1-CDK2 binding for this regulation.

The canonical *Cdk2ap1* gene structure is highly conserved across mammals (Fig. 4A and 4B). The predicted Cdk2ap1 ORFs from mouse, human, rhesus monkey, macaque, cattle, goat, pig and opossum genomes exhibit 86.1% identity at the amino acid level, all utilizing a conserved ATG start codon within the canonical exon 1 (fig. S4A). The alternative MT2B2 promoter for *Cdk2ap1* only exists in the mouse genome, and is absent in primate, livestock and rat genomes (Fig. 4B and fig. S4B). However, in all eutherian mammals examined, *Cdk2ap1*^*ΔN*^ are generated by species-specific promoters, all of which drive a transcript that skips exon 1 and directly splices into the canonical exon 2. All annotated Cdk2ap1^ΔN^ proteins utilize a conserved ATG start codon within the canonical exon 2 (Fig. 4B). In human preimplantation embryos, the predominant *CDK2AP1*^*ΔN*^ isoform is driven by a putative promoter region that contains an annotated L2a retrotransposon and a Charlie4z DNA transposon (Fig. 4A). Published ChIP-seq data in human ESCs supports *bona fide* promoter activity at the L2a/Charlife4z region, which exhibited an enrichment for H3K4Me3(*61*), H3K27Ac(*62*) and RNA polymerase II association (*63*) (fig. S4C). In human, rhesus monkey and marmoset genomes, orthologous *Cdk2ap1*^*ΔN*^ isoforms initiate transcription from this highly conserved L2a/Charlie4z region (fig. S4B), which contains predicted core promoter motifs, including an initiator motif near the TSS and a downstream DPE (downstream promoter element) motif (fig. S4B)(*64, 65*). Hence, the L2a/Charlie4z region likely possesses promoter activity to drive *CDK2AP1*^*ΔN*^ in multiple primates.

**Fig. 4.**
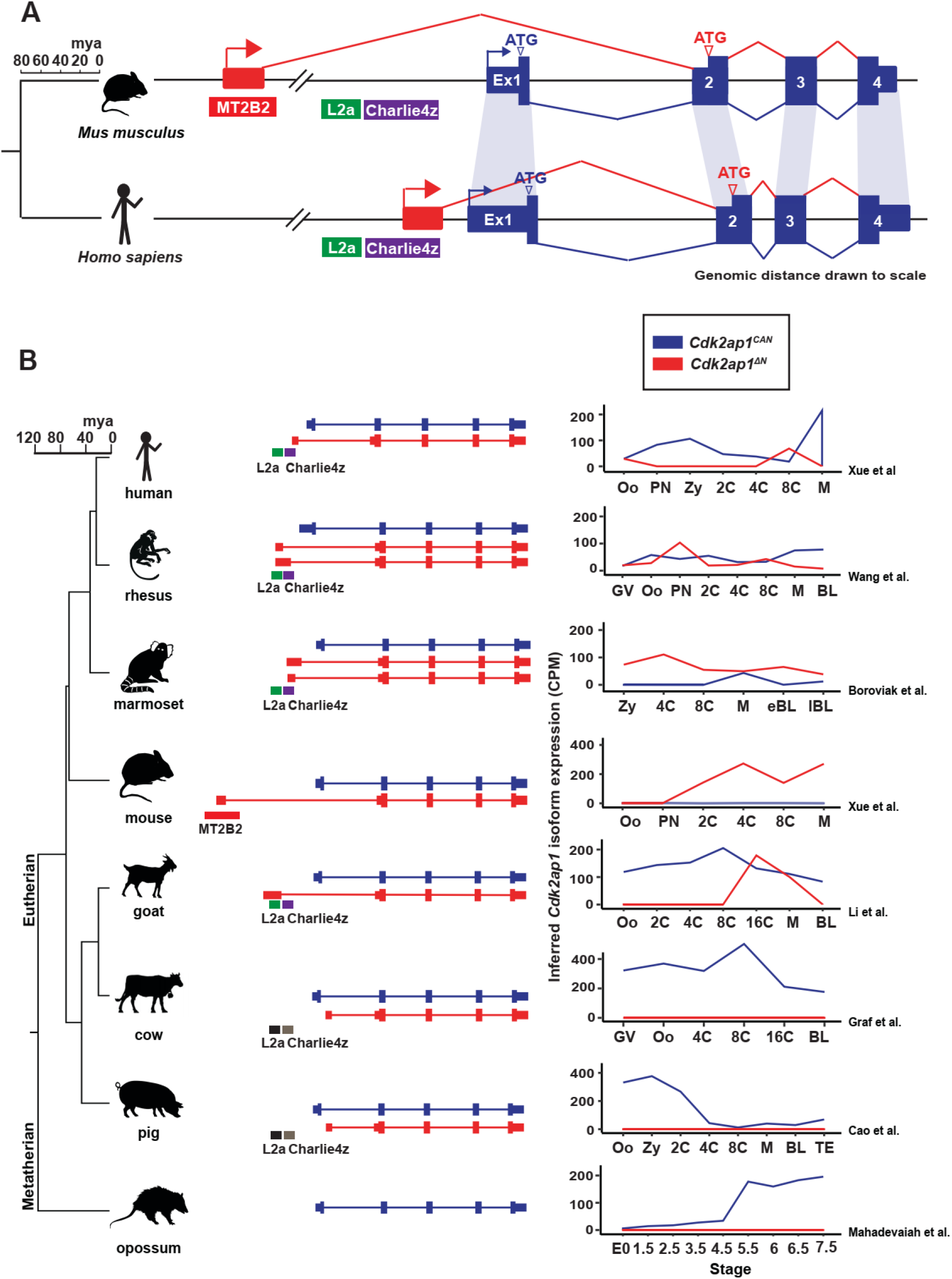
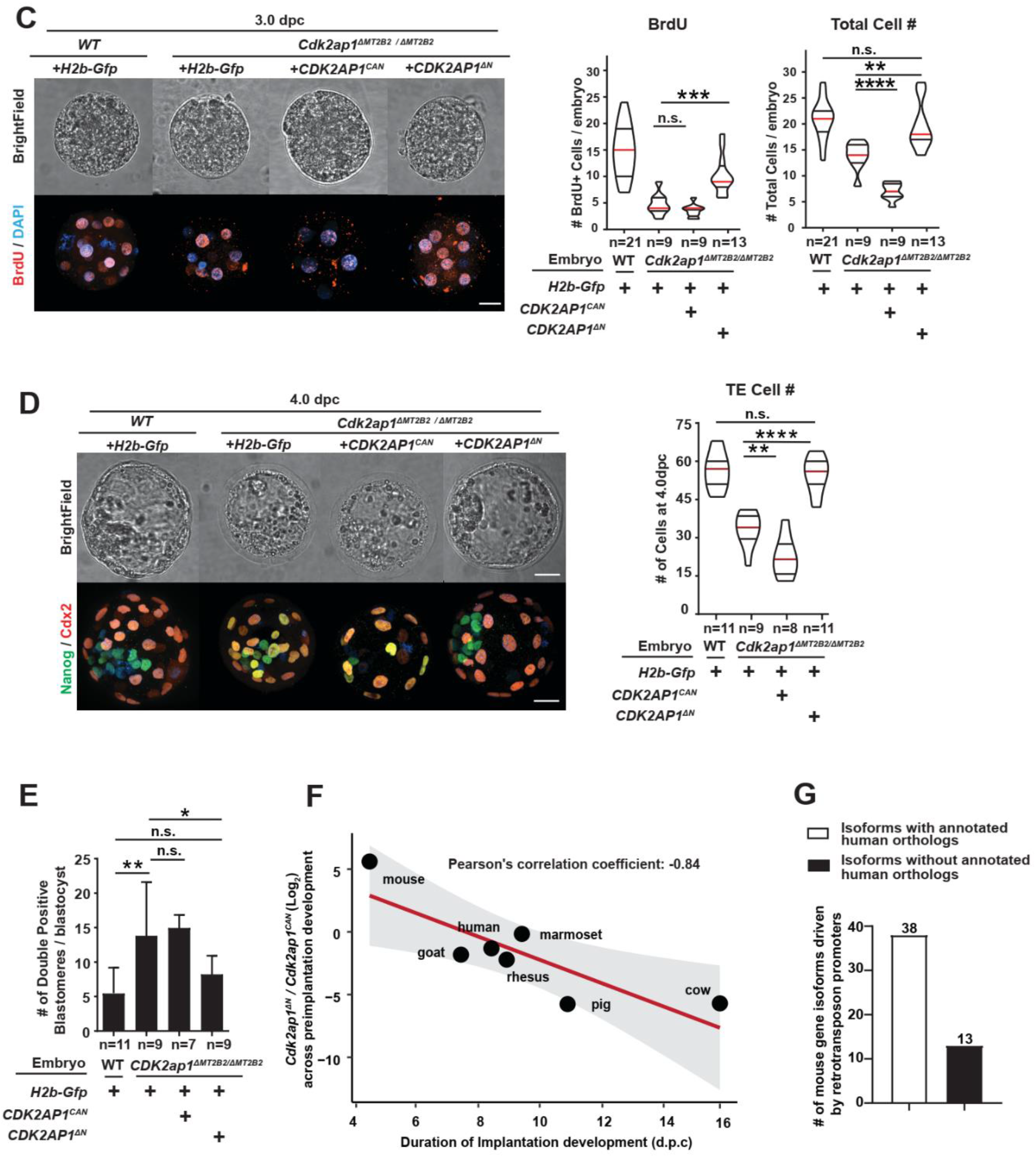
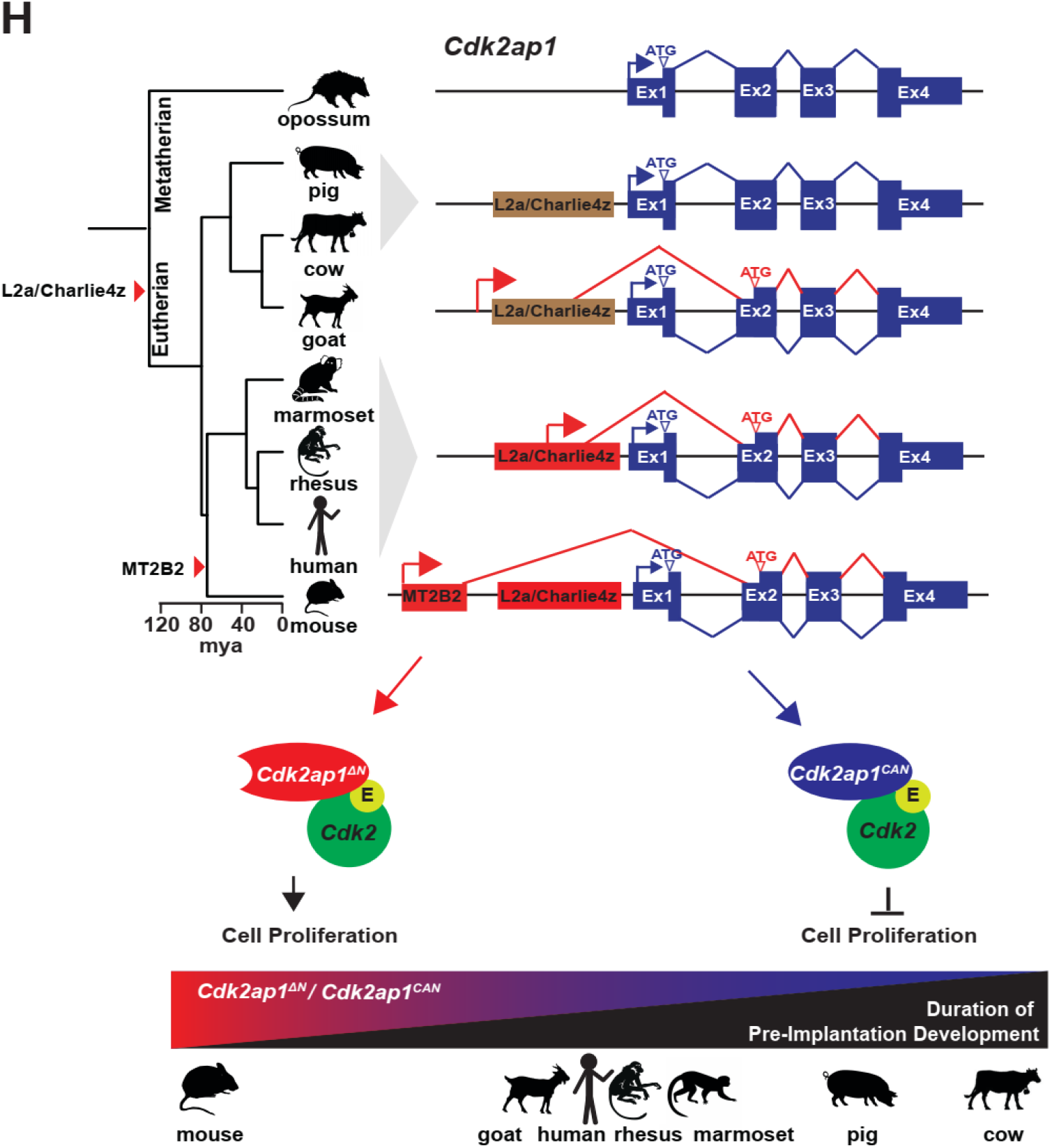
The MT2B2 driven, N-terminally truncated mouse Cdk2ap1 protein is evolutionarily conserved. **A**. The preimplantation-specific *Cdk2ap1*^*ΔN*^ isoforms are derived from species specific promoters in mouse and human but are highly conserved in protein-coding sequences. In preimplantation embryos, the mouse *Cdk2ap1*^*ΔN*^ isoform originates from the MT2B2 retrotransposon promoter; the human *CDK2AP1*^*ΔN*^ isoform originates from a genomic region containing an L2a element and a Charlie4z hAT DNA transposon element. Blue, canonical exons; red, alternative exons in preimplantation embryos. **B**. Canonical *Cdk2ap1* and N-terminally truncated *Cdk2ap1*^*ΔN*^ exhibit species specific differential expression in mammalian preimplantation embryos. Bioinformatics analyses on published RNA-seq data of mammalian preimplantation embryos revealed species specific regulation of *Cdk2ap1* and *Cdk2ap1*^*ΔN*^ (Sup Table S3). Isoform specific expression of *Cdk2ap1* in each species was determined by the total *Cdk2ap1* expression and the ratio between isoform specific splicing junctions. **C, D**. Cdk2ap1^*ΔN*^ is functionally conserved between human and mouse. Ectopic expression of *CDK2AP1*^*ΔN*^, but not *CDK2AP1*^*CAN*^, rescues the cell proliferation defects in *Cdk2ap1*^*ΔMT2B2/ΔMT2B2*^ morulae (**C**) and blastocysts (**D**), as demonstrated by BrdU incorporation and total cell number. **C**. Representative confocal images of BrdU staining (left) and quantification of BrdU incorporation and total cell number (right) were shown for 3.0 dpc embryos. **D**. Representative confocal images of Nanog and Cdx2 staining (left) and quantification of TE cell numbers (right) were shown for 4.0 dpc embryos. **C. D**. Scale bars 20 μm. Quantitation was shown as violin plots with median (red), lower (25%) and upper (75%) quartiles (black lines). **C**. *Cdk2ap1*^*ΔMT2B2/ΔMT2B2*^ *H2b-Gfp* vs *CDK2AP1*^*ΔN*^ (BrdU), *** *P* =0.0005, t=4.1, df=20; *Cdk2ap1*^*ΔMT2B2/ΔMT2B2*^ *H2b-Gfp* vs *CDK2AP1*^*CAN*^ (total cell number), \*\*\*\**P* < 0.0001, t=8.4, df=16; *Cdk2ap1*^*ΔMT2B2/ΔMT2B2*^ *H2b-Gfp* vs *CDK2AP1*^*ΔN*^ (total cell number), \*\**P* = 0.0031, t=3.4, df=20. **D**. *Cdk2ap1*^*ΔMT2B2/ΔMT2B2*^ *H2b-Gfp* vs *CDK2AP1*^*ΔN*^, \*\*\*\**P* < 0.0001, t=6.9, df=18; *Cdk2ap1*^*ΔMT2B2/ΔMT2B2*^ *H2b-Gfp* vs *CDK2AP1*^*CAN*^, \*\**P* = 0.007, t=3.1, df=15. **E**. Quantitation of Nanog and Cdx2 double positive cells in *Cdk2ap1*^*ΔMT2B2/ΔMT2B2*^ embryos overexpressing *CDK2AP1*^*ΔN*^ or *CDK2AP1*^*CAN*^. *H2b-Gfp*-overexpressing wildtype vs. *Cdk2ap1*^*ΔMT2B2/ΔMT2B2*^ embryos, ** *P* = 0.007, t=3.1, df=17; *H2b-Gfp* vs *CDK2AP1*^*ΔN*^ overexpression in *Cdk2ap1*^*ΔMT2B2/ΔMT2B2)*^ embryos, * *P* =0.04, t=2.3, df=18. **F**. The *Cdk2ap1*^*ΔN*^ to *Cdk2ap1*^*CAN*^ ratio is correlated with the duration of preimplantation development in multiple mammalian species. The *Cdk2ap1*^*ΔN*^ to *Cdk2ap1*^*CAN*^ ratio, calculated based on the sum of normalized RNAseq reads across isoform-specific junctions during preimplantation stages, is plotted against the average duration of preimplantation development for each species. Pearson’s correlation coefficient between Log_2_ (*Cdk2ap1*^*ΔN*^ */ Cdk2ap1*^*CAN*^) and duration of preimplantation development equals to −0.84, ** *P* = 0.018, t = −3.5, df = 5; the *P* value was calculated as part of the Pearson’s product-moment correlation. **G**. A subset of mouse specific retrotransposon promoters drive gene isoforms encoding evolutionarily conserved, N-terminally altered ORFs. Manual curation of the top 93 highly and differentially expressed retrotransposon promoters reveals 51 retrotransposon:gene isoforms, whose ORFs differ from those of the canonical isoforms. Among these 51 retrotransposon:gene isoforms with mouse retrotransposon promoters, 13 (26%) gene isoforms are orthologous to Refseq or Ensemble annotated human gene isoform. **H**. A diagram illustrates a model on the gene regulation on *Cdk2ap1* in mammalian preimplantation embryos. The canonical *Cdk2ap1* and *Cdk2ap1*^*ΔN*^ are two major *Cdk2ap1* isoforms in mammalian preimplantation embryos, which exhibit opposite regulation on cell proliferation. While the N-terminally truncated Cdk2ap1^ΔN^ isoform is highly conserved across multiple mammals, species specific regulation of *Cdk2ap1*^*ΔN*^ results in a varying degree of expression in preimplantation embryos. All 7 eutherian mammals examined contain the integration of an L2a element and a Charlie4z element upstream of *Cdk2ap1*. The transcription of *Cdk2ap1*^*ΔN*^ in preimplantation embryos originates from the L2a/Charlie4z region in human, rhesus monkey, marmoset and goat, but not in mouse, pig and cattle. Both canonical *Cdk2ap1* and *Cdk2ap1*^*ΔN*^ are expressed in human, rhesus monkey, marmoset and goat preimplantation embryos, but they peak at different developmental stages. A mouse-specific MT2B2 retrotransposon promoter promotes strong *Cdk2ap1*^*ΔN*^ expression in preimplantation embryos, making it the predominant *Cdk2ap1* isoform that is functionally essential. Pig and cattle exhibit sequence variations in the L2a/Charlie4z region compared to primates, and only express canonical *Cdk2ap1*, but not *Cdk2ap1*^*ΔN*^, in preimplantation embryos. Intriguingly, our data implicate that the *Cdk2ap1*^*ΔN*^ to *Cdk2ap1* ratio is inversely correlated with the duration of preimplantation development. All *P* values are calculated using unpaired two-tailed Student’s *t* test. n.s., not significant.

The human CDK2AP1^*Δ*N^ and mouse Cdk2ap1^*Δ*N(MT2B2)^ proteins share 97% sequence identity and exhibit functional conservation. Upon overexpression in mouse *Cdk2ap1*^*ΔMT2B2/ΔMT2B2*^ embryos, the human *CDK2AP1*^*ΔN*^ isoform mimicked the mouse *Cdk2ap1*^*ΔN(MT2B2)*^ isoform, restoring cell proliferation and cell fate specification to wildtype levels (Fig. 4C-4E). In contrast, the canonical human *CDK2AP1*^*CAN*^ isoform mimicked the mouse *Cdk2ap1* ^*CAN*^ isoform in *Cdk2ap1*^*ΔMT2B2/ΔMT2B2*^ embryos, further reducing BrdU incorporation and total cell number, particularly in the TE compartment (Fig. 4C and 4D). Hence, the opposite functions of *Cdk2ap1*^*CAN*^ and *Cdk2ap1*^*ΔN*^ is evolutionarily conserved.

The Cdk2ap1^*Δ*N^ isoform exhibits a spectrum of species-specific regulation modalities in mammalian preimplantation embryos. Human, rhesus monkey, marmoset and goat preimplantation embryos express both *Cdk2ap1*^*CAN*^ and *Cdk2ap1*^*ΔN*^ with different developmental stages for their peak expression; pig and cattle preimplantation embryos only express the canonical *Cdk2ap1;* mouse preimplantation embryos are characterized by the predominant *Cdk2ap1*^*ΔN(MT2B2)*^ expression (Fig. 4B, table S6).

The species-specific *Cdk2ap1*^*ΔN*^ regulation is achieved by its divergent promoters. The L2a/Charlie4z region is present in all 7 eutherian mammals examined (Fig. 4A, 4B). In human, and possibly other primates, the putative promoter in the L2a/Charlie4z region drives the transcription of *Cdk2ap1*^*ΔN*^ in preimplantation embryos (Fig.4B, fig. S4B). In pig and cattle, the L2a/Charlie4z region lacks promoter activity in preimplantation embryos, possibly due to sequence variations (Fig. 4B, fig. S4B). A more potent *Cdk2ap1*^*ΔN*^ induction occurs in mouse, where a recent MT2B2 integration generates a promoter, replacing the L2a/Charlie4z region as the key *Cdk2ap1*^*ΔN*^ promoter in preimplantation embryos.

Different mammals exhibit considerable phenotypical differences in preimplantation development. The duration of mammalian preimplantation development is highly variable between species, with 4.5 days for mouse and ≥10 days for cattle and pigs (fig. S4E and table S7). Intriguingly, the ratio of *Cdk2ap1*^*ΔN*^ to *Cdk2ap1*^*CAN*^ in preimplantation embryos is inversely correlated with the duration of preimplantation development across all 7 eutherian mammals examined (Fig. 4F). Mammalian blastocysts consist of 100-200 cells, and their competency for implantation roughly correlates with the absolute number of blastomeres during uterine apposition(*66*). Hence, we speculate that a higher abundance of Cdk2ap1^*Δ*N^ could serve to promote cell proliferation to reach competency for implantation sooner.

In addition to MT2B2, other mouse-specific retrotransposon promoters also drive gene isoforms that encode evolutionarily conserved, N-terminally altered ORFs (fig S4E). Among the top 93 most highly and differentially expressed mouse retrotransposon promoters, 14% (13/93) drive retrotransposon:gene isoforms that not only encode an altered ORF, but also have RefSeq/Ensemble annotated, orthologous human gene isoforms (Fig. 4G, fig. S4E and table S8). The protein sequence conservation of these orthologous gene isoforms spans ∼85 million years of human-mouse divergence, likely an indication of functional significance. The expression regulation of these orthologous gene isoforms is often divergent among species, and retrotransposons contributes to a key mechanism for species-specific gene regulation. Thus, the intricate interaction between retrotransposon promoters and host genomes contribute to species-specific gene regulation of evolutionarily conserved protein isoforms, yielding distinct expression patterns, important developmental functions and phenotypical diversity among species.

## Discussion

Colonization of transposable elements pose considerable threats to the integrity of mammalian genomes(*67, 68*), as they increase the risk of insertional mutagenesis(*69*–*71*), non-homologous recombination(*70*), and genome instability(*72, 73*). However, transposable elements also provide abundant genetic material for gene regulatory sequences, substantially increasing the species-specific complexity of gene regulation and transcript diversity, ultimately driving genome innovation(*7, 16*). Deletions of specific retrotransposon-derived gene regulatory elements can alter the structure and expression of proximal host genes(*15, 21, 74, 75*), yet it is unclear to what extent such transposable elements regulate important biology in the host.

To date, the best characterized gene regulatory transposable elements are retrotransposon promoters that drive species-specific gene isoforms to regulate fertility or enrich phenotypical variance, but they are often non-essential for viability(*25, 58*). For example, an intronic mouse MTC promoter drives an N-terminally truncated, oocyte specific Dicer^O^ isoform with enhanced

Dicer activity to safeguard meiotic spindle formation(*25*). An intronic *Shellder* element in *Drosophila simulans* alters the splicing of the calcium-activated potassium channel *Slo*, causing natural variation of courtship songs(*58*). To our knowledge, the mouse MT2B2 element upstream of *Cdk2ap1* is among the first essential retrotransposon promoters characterized in mammals. The MT2B2 retrotransposon drives an N-terminally truncated *Cdk2ap1*^*ΔN*^ isoform that promotes cell proliferation, controls blastocyst cell numbers, and ultimately, regulates embryo implantation. The cell proliferative function of *Cdk2ap1*^*ΔN*^ is opposite from the anti-proliferative function of canonical *Cdk2ap1*^*CAN*^, both exerting their effects on Cdk2. It is possible that *Cdk2ap1*^*ΔN*^ and *Cdk2ap1*^*CAN*^ also regulate additional cell proliferation pathways(*53, 76, 77*), because *Cdk2* knockout alone is not sufficient to render preimplantation defects (*78*). While *Cdk2ap1*^*ΔN*^ and *Cdk2ap1*^*CAN*^ are both essential for mouse development, their distinct induction in preimplantation and mid-gestation embryos, respectively, dictates the specific biology they each regulate.

The N-terminally truncated Cdk2ap1^*Δ*N^ proteins are present in many mammalian species, exhibiting a highly conserved protein sequence and cellular function. In comparison, the gene regulatory mechanisms of Cdk2ap1^*Δ*N^ are divergent in mammals, as species-specific transcriptional regulation governs the expression of these orthologous Cdk2ap1^*Δ*N^ isoforms. While ancient transposable elements could be present in mammalian genomes, most transposon integrations are unique events in the evolutionary history of a species. A subset of these transposable elements play an important role in the evolution of gene regulation by enabling the rapid emergence of regulatory elements.

In all eutherian mammals examined, the ancient integration of an L2a element and a Charlie4z element is present upstream of *Cdk2ap1*. The L2a/Charlie4z region possesses promote activity to drive modest *Cdk2ap1*^*ΔN*^ induction in human, and possibly in other primates. Sequence variations within the L2a/Charlie4z element are correlated with the lack of *Cdk2ap1*^*ΔN*^ promoter activity and absence of *Cdk2ap1*^*ΔN*^ expression in pig and cattle. Interestingly, a mouse-specific MT2B2 integration yields a second *Cdk2ap1*^*ΔN*^ promoter, driving its strong induction in preimplantation embryos as an essential gene. It is unlikely that the MT2B2 element is essential for development at the time of its integration. The strong *Cdk2ap1*^*ΔN*^ induction driven by MT2B2 may induce additional gene regulatory changes during evolution, and eventually rendering it indispensable for preimplantation development. This transposon-dependent *Cdk2ap1*^*ΔN*^ regulation yields species-specific difference in *Cdk2ap1*^*ΔN*^ abundance in preimplantation embryos. Intriguingly, the relative *Cdk2ap1*^*ΔN*^ abundance in each examined species is inversely correlated with the duration of preimplantation development. This is consistent with the idea that biological differences among species often stem from different gene regulatory mechanisms, rather than different protein sequences(*79*). Taken together, transposable elements can yield diverse gene regulation of an evolutionarily conserved protein isoform, orchestrate species-specific expression and developmental functions, and may eventually evolve to be essential.

Unlike most somatic tissues, mammalian preimplantation embryos are unusually permissive to retrotransposon induction. Numerous retrotransposon elements generate preimplantation-specific gene isoforms, characterized by alternative transcriptional/translational regulation, and in some cases, new ORFs and novel protein functions. Hence, retrotransposons are important building blocks for evolutionary “tinkering”, promoting species-specific gene innovation and possessing the capacity to generate functionally essential protein isoforms.

## Supporting information

DropBox Link to Table S1, S2 and S3

## Acknowledgments

We thank Dr. P Lishko and H. Dhaliwal for advice and assistance on embryo culture and embryo manipulation, Dr. J. Cox for confocal microscope access, Dr. S. Dey for advice on uterine and implantation phenotype, Dr. Y. Zhou and Dr. H. Huang for bioinformatics assistance, Dr. M. Rape, Dr. A. Manford and Dr. F. Rodriquez for advice and assistance on protein purification, H. Nolla and K. Heydari for FACS training, M. West for advanced imaging training, H. Asahara for sequencing and cloning assistance, A. Killilea for cell culture advice and reagents, E. Newman, G. Wang, E. Tang and W.C. Chen for managing mouse colonies, R. Medina and A. Castillejo for lab maintenance. We appreciate stimulating discussions with Dr. R Rogers, Dr. M Slatkin, Dr. M. Nachman and Dr. S Banker on evolutionary biology, and scientific support from the stem cell center, particularly Lily Mirels. Special thanks to R. Valbuena for critical experimental assistance throughout the project, Dr. D. Yi for microinjection comparison, C. DiBiaggio for testing and optimizing guide RNA conditions, M. Kinisu and Dr. S. Chen for proofreading and revising the manuscript, as well as all members of the He lab for discussion of the project.

## Author contributions

A.J.M, D.R. and L.H. conceived the key hypotheses of this study and designed the majority of the experiments. A.J.M. established mouse embryo collection, culture and manipulation systems, and developed CRISPR-EZ to generate deletions, epitope tagging and electroporation-based overexpression experiments in mouse embryos. A.J.M phenotypically characterized all mouse mutants, discovered the opposite functionality of two Cdk2ap1 isoforms, and investigated orthologous human Cdk2 isoforms. A.L. conducted the embryo transfer experiments to generate the edited animals in this study. F.X. characterized implantation defects of *Cdk2ap1* mutants. K.T. performed mouse husbandry, genotyping, cloning and vector generation, and collected post implantation embryos for expression analysis to determine peak Cdk2ap1 expression. D.R, T.W. and T.S conceived the bioinformatics pipeline to analyze the expression of retrotransposon family and retrotransposon:gene isoforms. W.S. and D.R. established the computational algorithms for analyzing and quantifying retrotransposon expression and retrotransposon:gene isoforms using RNA-seq data, and analyzed all RNA-seq datasets for mammalian preimplantation embryos.

A.J.M. performed literature search to identify all RNA-seq datasets for preimplantation embryos for our bioinformatic analyses. J.C. and A.B. polished RNA-Seq mapping and generated expression and lists of retrotransposon:gene isoforms in mouse and human. M.N. performed manual curation of 250 retrotransposon:gene isoforms to generate supplementary table S5. G.S. performed bioinformatic analysis to characterize the ORFs encoded by the retrotrotransposon:gene isoforms. D.S. T.W. and L.H provided guidance to the key experiments and bioinformatics analyses. A.J.M and W.S. generated all figure panels and most supplementary tables. A.J.M and L.H. drafted and revised the manuscript, all other co-authors proof-read the manuscript.

## Data and materials availability

All data is available in the main text or the supplementary materials. All data and reagents will be available to any researcher for purposes of reproducing or extending the analysis.

## Inclusion and diversity statement

We worked to ensure gender balance in the authors who contributed to this paper

## Supplementary Materials

Materials and Methods

Figures S1-S4

Tables S1-S9

References (80-96)

