## Supplemental Material for "A species-specific retrotransposon drives a conserved *Cdk2ap1* isoform essential for preimplantation development"

**This PDF file includes:**

Materials and Methods
Supplementary Text
Figs. S1 to S4
Tables S1 to S9

#### Materials and Methods

##### Bioinformatic analyses

*RNA-seq data processing*

RNA-seq raw sequencing files for mammalian preimplantation embryos were downloaded from NCBI Sequence Read Archive and EMBL-EBI ArrayExpress (Supplementary Table S1). After trimming off adapter sequences with cutadapt (v. 2.10)(*1*), RNA-seq reads were mapped to the reference genomes using STAR (v. 2.7.1a)(*2*). To increase the detection sensitivity of spliced RNA-seq reads, we applied the two-pass alignment strategy(*3*). For the first pass alignment, we aligned RNA-seq reads using STAR genome index files generated with the gene annotations provided by RefSeq (Supplementary Table S1). Subsequently, we collected all the detected splice sites for each mammalian species and updated the STAR genome index files by incorporating previously unannotated splice sites. To ensure the accuracy of the updated STAR index, we only considered splice sites that were confirmed by at least 3 mapped reads and were characterized by STAR-defined canonical intron motifs. These updated STAR genome index files were then employed for the second round of sequence alignment. To further reduce the number of spurious junctions, we only kept reads containing junctions that were included in the SJ.out.tab files (STAR option: --outFilterType BySJout). All the raw RNA-seq sequencing data used in this study are available from the Gene Expression Omnibus, ArrayExpress, or Short Read Archive, at accessions GSE44183, GSE45719, GSE36552, GSE86938, E-MTAB-7078, SRA076823, GSE139512, GSE143850, GSE52415, GSE129742, E-MTAB-7515.

*Expression quantification of protein coding genes and retrotransposons*

To obtain genomic coordinates of protein-coding genes and non-coding transcripts, we processed the gene annotation files provided by Refseq (Supplementary Table S1). To obtain genomic coordinates of retrotransposon families, we downloaded the Repeatmasker output from UCSC and NCBI and selected for elements that belong to LINE, SINE and LTR. To avoid confounding between gene and retrotransposon expression, we excluded all the retrotransposons that overlap with Refseq annotated gene exons from our retrotransposon quantitation. We then quantified gene and retrotransposon expression by counting uniquely mapped RNA-seq reads that overlap with annotated exons or retrotransposons using FeatureCounts(*4*) (v. 1.6.3, options -O -B -p --fracOverlap 0.1 -M --fraction -T 5 -Q 255 for paired end RNA-seq samples, -O --fracOverlap 0.1 -M --fraction -T 5 -Q 255 for single end RNA-seq samples). The number of reads mapped to all the members of a retrotransposon subfamily were then combined to obtain retrotransposon subfamily-level expression. Although our expression quantification strategy doesn't account for multiply mapped reads and thus likely underestimates the expression of retrotransposons, it is the safest strategy to accurately evaluate retrotransposon expression.

*Differential expression analysis on protein coding genes and retrotransposons*

We combined the expression value of protein-coding genes and retrotransposon subfamilies into a single matrix and retained only the genes or the retrotransposon subfamilies with at least one CPM (counts per million) in at least one sample. For datasets with more than 2 samples per developmental stage, we used edgeR (v.3.12.0)(*5*) to test for differential expression during preimplantation development (negative binomial likelihood ratio test after full-quartile normalization(*6*) and RUVr normalization(*7*)). Genes or retrotransposon subfamilies with an FDR (false discovery rate) less than 0.05 were defined as differentially expressed (DE). For datasets with only one sample per developmental stage, we inferred the degree of differential expression by calculating the standard deviation per gene using its expression values across all developmental stages. All expressed protein coding genes or retrotransposon subfamilies were then ranked by averaged expression signal during the peak developmental stage (Supplementary Table S1, S2). To illustrate the dynamic expression of protein-coding genes and retrotransposons, we generated heatmaps using z-scores of top highly and differentially expressed protein-coding genes or retrotransposon subfamilies. Z-score was defined as the standard deviations by which the expression value of a gene or a retrotransposon subfamily is above or below its mean expression across all the preimplantation stages. Hierarchical clustering was then performed to group genes or retrotransposon subfamilies with similar expression patterns together. Extreme z-scores (below 0.01 quantile or above 0.99 quantile) were capped for display purposes.

*Annotation of retrotransposon:gene junction reads*
We first performed transcript assembly using StringTie2 to identify novel exon structures that were absent from Refseq annotation(*8*). We then extracted split RNA-seq reads from aligned .bam files and only kept reads that have at least 6 nucleotides mapped to the genome at both ends. Only reads with splicing junctions between 50 and 100,000 bp in length in the genome were retained. A read was considered as a retrotransposon:gene junction read when it fulfilled the following two criteria: 1) both ends of the read were mapped to exons (assembled exons from RNA-seq data or annotated exons from RefSeq); 2) one end of the read was mapped to annotated protein-coding gene exons and the other end was mapped to an annotated retrotransposon. We then counted the number of retrotransposon:gene junction reads for each unique splicing junction. Due to the repetitive nature of retrotransposon sequences, this procedure may not be entirely accurate, especially in the presence of gene families and/or pseudogenes. Hence, only junctions with at least 10 reads in at least one samples were retained for downstream differential expression analysis.

*Differential expression of retrotransposon:gene junction reads:*
For datasets with more than 2 samples per developmental stage, we used edgeR (v.3.12.0)(*5*) to test for differential expression of retrotransposon:gene junction reads during preimplantation development. Negative binomial likelihood ratio test was performed after full-quartile normalization(*6*) and RUVr normalization(*7*). Junctions with an FDR (false discovery rate) less than 0.05 were defined as differentially expressed (DE). For datasets with only one sample per developmental stage, we inferred the degree of differential expression by calculating the standard deviation per gene using its expression values across all developmental stages.

*Manual annotation of the gene structure and ORF conservation of retrotransposon:gene isoforms*
Following the bioinformatic identification of retrotransposon:gene isoforms, a manual annotation step was performed to define their gene structures and ORFs. The top 250 most differentially expressed mouse retrotransposon:gene junctions were selected for manual curation, and the position of retrotransposon element within the retrotransposon:gene isoforms were classified as 5', internal, or 3'. 5', the retrotransposon element is transcribed at the 5' end of a retrotransposon:gene isoform, and it only splices into a downstream host gene exon. Often time such retrotransposon elements act as promoters that harbor transcriptional start sites (TSS). Internal, the retrotransposon element contributes to an internal exon of a retrotransposon:gene isoform, with splicing to both upstream and downstream host gene exons. Such retrotransposons are exonized into host genes. 3', the retrotransposon element is transcribed at the 3' end of a retrotransposon:gene isoform, as

we only detect splicing between retrotransposons and upstream gene exons. These retrotransposons likely provide polyadenylation signals.

*Differential expression of Cdk2ap1 isoforms:*

For *Cdk2ap1* isoform expression analyses, we first combined RefSeq annotation for *Cdk2ap1* with our transcript assembly results to obtain a more comprehensive catalog of *Cdk2ap1* gene isoforms in each species. While multiple Cdk2ap1 isoforms exist for a given species, these isoforms encode either the canonical Cdk2ap1 protein or N-terminally truncated Cdk2ap1 isoforms. To infer *Cdk2ap1* isoform-level expression, we redistributed *Cdk2ap1* gene-level RNA-seq signal to canonical and N-terminally truncated isoforms based on the amount of RNA-seq reads across isoform specific splicing junctions.

From the top 250 most highly and differentially expressed mouse retrotransposon:gene isoforms, we selected all that were driven by retrotransposon promoters for manual annotation of ORFs (n=93). In parallel, we also selected the top 100 from the top 250 most highly and differentially expressed mouse retrotransposon:gene isoforms for ORF annotation. Retrotransposon:gene isoforms were reconstructed into a predicted mRNA based on RNA-seq data and mouse reference genome assembly mm10. The ORF of each retrotransposon:gene isoform was predicted from the reconstructed mRNA using SnapGene 2.3.2., and then compared to the ORF encoded by the most highly-expressed isoform among the remaining gene isoforms in preimplantation embryos. Interestingly, a small subset of retrotransposon:gene isoforms, mostly internal cases, constitute 100% preimplantation specific transcripts of the corresponding gene, and exhibit no ORF alterations. The predicted ORF alterations were classified into 6 categories: deletion, insertion, replacement, other N-terminal modifications, N.A. (no modification to protein) and N.D. (not determined). For each of the mouse retrotransposon:gene isoforms encoding an altered ORF, we examined the human RefSeq and Ensemble annotations to identify the annotated human gene isoform that encodes the orthologous ORF. All analyses were done using the mouse reference genome assembly mm10 and the human reference genome assembly hg38.

*ENCODE and Roadmap Epigenomics project data analysis and CDK2AP1 promoter activity.*

The wiggle files for ChIP-seq of H3K4me3(*9*) (GSM733657), H3K27Ac(*10*) (GSM646336), and Pol2(*11*) (GSM748532) were obtained from Cistrome database(*12*) and displayed using UCSC genome browser(*13*). The human H1-ESC RNA-seq data from GSE23316(*9*) were downloaded from NCBI GEO database(*14*), and Kallisto(*15*) was used to quantify the isoform expression levels with GENCODE (GRCh38 ver. 26)(*16*). Four RNA-seq replicates with insert length of 200bp were used.

*Phylogenetic analysis.*

Genomic Phylogeny of various placental mammal taxa were generated by first organizing a selection of animal of interest in terms of their binomial nomenclature in Latin. This list is then imputed into TimeTree.org(*17*), which generates timescales and species divergence nodes as a Newick file. This file is imported and modified using FigTree v1.4.4 for presentation purposes.

*Sequence Alignment*

Current Sequence alignment was performed using clustal Omega (https://www.ebi.ac.uk/Tools/msa/clustalo/) with default parameters. Alignment files were used as input for alignment shading (BoxShade v3.21 https://embnet.vital-it.ch/software/BOX_form.html). Protein alignments utilized peptide sequences from each species provided by NCBI GenPept entries. Genomic sequence alignments for L2a and Charlie4z utilized the DNA sequences around L2a/Charlie4z regions provided by UCSC. Zoomed in alignment for Charlie4z was manually adjusted and annotated for core promoter elements.

**Mouse embryo culture, engineering, molecular analyses and phenotypical analyses.**

*Mouse Embryo Isolation and Culture*

Three-to-five-week-old C57BL/6J female mice (Jackson Laboratory, 000664) were superovulated by intraperitoneal (IP) injection of 5 IU of Pregnant Mare Serum Gonadotropin (PMSG, Calbiochem, 367222), and 46-48 hours later, 5 IU of Human Chorion Gonadotropin (hCG, Calbiochem, 230734). Superovulated females were each housed at a 1:1 ratio with a 3-8-month-old C57BL/6J male to generate 1-cell zygotes at 0.5 days post coitum. Using forceps under a stereomicroscope (Nikon SMZ-U), the ampulla of oviduct was nicked, releasing fertilized zygotes associated with surrounding cumulus cells into 50 µl M2 + BSA media (M2 media (Millipore, MR-015-D) supplemented with 4 mg/mL bovine serum albumin (BSA, Sigma, A3311)). Using a handheld pipette set to 50 µl, we dissociate zygotes from cumulus cells, after the cumulus oocyte complexes were incubated for 2 min in a 200 µl droplet of 1X Hyaluronidase in M2 solution (Millipore, MR-051-F), followed by five washes in the M2+BSA media to remove cumulus cells. From this point on, embryos were manipulated using a mouth-controlled assembly consisting of a capillary pulled from glass capillary tubes (Sigma, pack of 250: P0674) over an open flame attached to a 15-inch aspirator tube (Sigma, pack of 5, A5177). Embryos were then transferred to the KSOM + BSA media (KCl-enriched simplex optimization medium with amino acid supplement (Zenith Biotech, ZEKS-050), supplemented with 1 mg/ml BSA), which was equilibrated to final embryo culture conditions at least 3–4 hours prior to incubation to reach optimal temperature, $CO_2$ and pH conditions. Embryos were cultured in 30 µl droplets of KSOM + BSA overlaid with mineral oil (Millipore, ES-005-C) in 35 × 10 mm culture dishes (CellStar Greiner Bio-One, 627160) in a water-jacketed $CO_2$ incubator under hypoxic conditions (5% $O_2$, 5% $CO_2$, 37 °C and 95% humidity).

*Single-Embryo Quantitative Reverse Transcription PCR (qRT-qPCR)*

All single-embryo cDNA was prepared using a modified version of the Single Cell-to-Ct qRT-PCR kit (Life-Technologies, 4458236). Whole embryos were isolated at a desired developmental stage and passed through three PBS washes. With a hand-held pipette set to 1 µL, a single embryo was collected in PBS and transferred to one tube of an 8 well PCR strip, and the presence of embryo was visually confirmed under microscope. To account for the larger sample input, we incubated each embryo in 20 µl "Lysis/DNAse" reagent at room temperature (25$^{o}$C) for 15 minutes, then added 2 µl of "Stop Solution" for a 2 min incubation at room temperature. Half reaction was stored at -80$^{o}$C as a technical replicate, and the remaining sample (11 µl) continued through the Single Cell-to-Ct protocol per manufacturer's recommendation. For each experiment, a single embryo was collected and reserved as a "-RT" control, 1 µl of PBS was collected as a "No Template Control". All qRt-PCR analyses were performed on the StepOnePlus Real Time PCR system (Thermo, 437660). All real-time qPCR analyses were performed using SYBR FAST qPCR Master Mix (Kapa Biosystems, KK4604) following manufacturer's protocol. Real time PCR analyses on retrotransposons detect their expression at the family level, using primers designed from the retrotransposon consensus sequences. To detect retrotransposon gene isoform expression, primers were designed against the predicted isoform and to span the retrotransposon:gene splicing junctions, with one primer located within the retrotransposon sequence and the other located within the gene exon. *Rfx1* was used as a reference for both mRNA and retrotransposon quantitation in real time PCR analyses. All real time PCR primers used in our studies are listed in Supplementary Table S9.

*Validation of Retrotransposon Gene Junction Reads by TA cloning*.

Upon completion of qRT-PCR analysis, the amplification samples were mixed at a 1-to-1 ratio with non-processive TAQ-polymerase supplied as a 2x Master Mix (Promega, M7123) and incubated at 72$^{o}$C for 10 min in order to append a single deoxyadenosine to the 3' ends of the amplicon. The amplified fragments that captured retrotransposon:gene junction reads were purified through gel extraction (BioBasic, BS654) before TA cloned into pGEM-T easy vector (Promega, A1360). The plasmids were sequenced by Sanger Sequencing at the UC Berkeley DNA Sequencing Facility, and the retrotransposon:gene junctions were analyzed and visualized using SnapGene (version 2.3.2).

*Rapid Extension of cDNA Ends (RACE)*

All RACE experiments were conducted following manufacturer's instructions (Clontech, 634858) with the following modifications. Input RNA was provided by pooling approximately 50 Morula stages mouse embryos followed by trizol RNA extraction per manufacturer's instruction (Life Technologies, 15596). A list of primers used in this experiment is listed in Supplementary Table S9.

*Luciferase Assay for translation efficiency*

To analyze the impact of retrotransposon derived 5'UTR on translation efficiency, we constructed luciferase reporters for translational assay using psiCheck2 luciferase reporter vector (Promega, C8021). The 5'UTRs of mouse canonical *Cdk2ap1* and *Cdk2ap1*$^{\Delta N}$ isoforms were cloned immediately upstream of the Renilla Luciferase ORF; the FireFly luciferase reporter cassette from the siCheck2 vector was cloned as a control. All reporters were *in vitro* transcribed, 5' capped and polyadenylated. *Renilla Luciferase* and *FireFly luciferase* reporter mRNAs were co-transfected into HEK293T cells (600 ng *Renilla Luciferase* mRNA and 2200 ng *FireFly luciferase* mRNA per well of a 12-well plate), using Lipofectamine 2000 (Life Technologies, 11668027). Approximately 8 hours later, samples were assayed for luciferase activity by Dual-Luciferase® Reporter Assay System (Promega, E1910) as per manufacturer's instructions using a Glomax 20/20 Luminometer (Promega).

*Mouse genome engineering by CRISPR-EZ*

Embryos were edited following the published CRISPR-EZ protocol(*18*, *19*). Briefly, super ovulated C57BL/6J female mice were used to generate pronuclear stage embryos. Pronuclear stage embryos were dissociated from cumulus cells using Hyaluronidase (Millipore, MR-051-F), the zona was weakened with acid Tyrode's solution (Sigma, T1788), and the embryos were subsequently washed in M2 buffer. For the MT2B2 deletion or *Cdk2ap1* exon 1 deletion, Cas9/sgRNA RNP complexes were assembled *in vitro* in a total of 10 μL by combining Cas9 protein (8 μM final concentration) with two sgRNAs (2 μg per sgRNA) flanking the desired deletion. Assembled RNPs were then mixed with 50-75 zygotes in 10 μL OptiMEM media (Thermo, 31985062), and total 20 μL mixture was delivered into zygotes by electroporation

(BioRad Genepulser XL, 1652660). Electroporation conditions were 30V, 6 Pulses, 3ms pulse length and 100ms Pulse interval. Electroporated embryos were immediately transferred into pseudo-pregnant CD-1 recipient females to generate genetically engineered mice. The *Cdk2ap1*$^{\Delta MT2B2/\Delta MT2B2}$ and *Cdk2ap1*$^{\Delta CAN/\Delta CAN}$ mice were maintained on an isogenic C57BL/6J background and housed in a non-barrier animal facility at UC-Berkeley.

For endogenous V5 tagging to *Cdk2ap1* isoforms, a synthesized donor oligo (IDT) for Homology Directed Repair (HDR) was added to the Cas9/sgRNA RNP Complex mixture at a final donor oligo concentration of 20 μM for CRISPR-EZ(*18*, *19*). Electroporated embryos were then cultured to appropriate developmental stages, fixed and processed for immunofluorescence staining (see below).

Correctly engineered mouse embryos or adult mice were confirmed by genotyping analyses. To extract DNA from embryos, embryos were washed twice with PBS, and 1 μl of PBS solution containing a single embryo was transferred into 10 μL of embryo lysis buffer (50 mM KCl (Fisher, catalog no. P217-3), 10 mM Tris-HCl, pH 8.5 (Fisher, BP1531), 2.5 mM $MgCl_2$ (Fisher, M33-500), 0.1 mg/ml gelatin (Fisher, G7–500), 0.45% Nonidet P-40 (Fluka, 74385), 0.45% Tween 20 (Sigma, P7949-500), and 0.2 mg/ml proteinase K (Fisher, BP1700-100)). Lysis was performed in a thermocycler with the following conditions: 55 °C for 4 h, 95 °C for 10 min, and 10 °C hold. At this point, 3-4 μl of the 11 μl of lysed material were used directly in a standard PCR reaction for genotyping. To extract DNA from mouse tails, we used a standard Proteinase K extraction protocol. All genotyping primers are listed in Supplementary table S9.

*mRNA Electroporation into mouse zygotes*

Conditions for mRNA electroporation were identical to the parameters described by the CRISPR-EZ protocol(*18*, *19*), except that mRNA was electroporated in place of RNP/sgRNA complexes. Prior to electroporation, *H2b-Gfp* control and *Cdk2ap1* mRNAs were prepared by *in vitro* transcription (IVT) using the Hiscribe T7 ARCA w/ Tailing kit following manufacturer's instructions (NEB, e2060). For each electroporation, 200 ng of control *H2b-Gfp* and 2000ng of experimental mRNA was mixed with 20 μl of Opti-MEM and combined with 25-75 mouse zygotes. Following electroporation, embryos were recovered and washed with M2 media and cultured under mineral oil in KSOM+BSA until the appropriate developmental stage for subsequent analyses. A list of IVT templates and primers was summarized in Supplementary table S9.

*Immunofluorescence staining on preimplantation embryos*

Embryos were fixed in 4% paraformaldehyde (Electron Microscopy Sciences, 19202) for 15 min at room temperature, and then transferred to wash buffer (PBS containing 0.1% bovine serum albumin, Sigma, A3311). Embryos were permeabilized with PBS containing 0.1% Triton X-100 and 0.1% BSA for 5 min, blocked for 1 hour at room temperature in PBS containing 10% goat serum (Fisher 31872) and 0.1% BSA, then incubated with appropriate primary antibody in blocking solution at 4 °C overnight. The primary antibodies include antibodies against Cdx2 (1:100, Abcam, ab76541), Nanog (1:100, CosmoBio, REC-RCAB0002PF), Cdk2ap1 (1:50, Santa Cruz sc-390283), V5 (1:100, a gift from the Tjian Lab), BrdU (1:100, Thermo Fisher, 17-5071-41). On the following day, embryos were passed through room temperature wash buffer wash buffer (PBS containing 0.1% bovine serum albumin, Sigma, A3311) twice before being incubated with appropriate secondary antibodies diluted in blocking solution at 4 °C overnight. The secondary antibodies used in our studies include goat anti-mouse IgG Alexa Fluor 594 (1:400, ThermoFisher, A11005), goat anti-rabbit IgG Alexa Fluor 594 (1:400, Thermo Fisher, A11037), goat anti-mouse IgG Alexa Fluor 488 (1:400, Thermo Fisher, A11001) and goat anti-rabbit IgG Alexa Fluor 488 (1:400, Thermo Fisher, A110034). Finally, embryos were stained with (4′,6-diamidino-2-phenylindole) (DAPI at 300 nM, Sigma, D9564) and subjected to imaging analyses using spinning disk scanning confocal microscopy (Nikon Eclipse TE200-E). After imaging, embryos were collected in the order they were imaged, lysed with temperature induced reverse-crosslinking and subjected to PCR based genotyping analysis.

*BrdU incorporation in preimplantation embryos*

Morulae and blastocysts were processed for BrdU analysis as previously described(*20*). Briefly, embryos were cultured for 1 hour in 20 µl droplet of KSOM + BSA media supplemented with 25 µM BrdU (BD Pharmingen, 51-2420KC) under mineral oil. Embryos were then washed three times in wash buffer (PBS containing 0.1% BSA, Sigma, A3311), then fixed in 4% paraformaldehyde for 10 min at room temperature. Embryos were washed again three times in wash buffer. Embryo permeabilization and DNA denaturation was performed simultaneously by incubating the embryos in 2M HCl/0.5% Triton-X100 in PBS for 20min (Triton X-100, Sigma, X100, HCl, Macron 2062-46). Embryos were washed again three times and placed in blocking solution (PBS containing 10% goat serum and 0.1% BSA) for 1 hour at room temperature. Embryos were incubated overnight at $4^{o}$C with anti-BrdU antibody in blocking buffer (1:100, Thermo, 17-5071-41), then processed for confocal imaging.

*Phenotypical analyses of embryo implantations*

For each deletion strain, uteri from littermate WT and *Cdk2ap1*$^{\Delta MT2B2/+}$ and littermate WT and *Cdk2ap1*$^{\Delta CAN/+}$ were collected from female mice at specific developmental stages for embryo implantation analyses. Implantation was considered abnormal if sites were spaced either shorter or further than the expected normalized inter-embryo distance. Collected uterus was cleared of attached fat tissue and photographed next to ruler for scaling and measurement purposes. Embryos were then surgically removed, small sample collected, washed twice in PBS and collected for PCR based genotyping analysis, as previously described above.

**Biochemical Analyses of Cdk2ap1 activity on cell proliferation.**

*Cell culture and transfection, Co-immunoprecipitation (Co-IP) and western analyses*

Transfection of HEK293T cells with pMSCV expression vectors for control (empty vector), *Cdk2ap1*$^{\Delta N(MT2B2)}$*-HA*, or *Cdk2ap1*$^{CAN}$*-HA* C-Terminally HA-tagged *Cdk2ap1* isoforms was performed using by standard polyethylenimine (PEI) transfection (Polysciences, 23966-1). For each, 10 µg of DNA were used for each 10cm dishes of HEK293T cells, where the ratio of DNA to PEI is 1:20. Transfected cells were collected at 48 hours, washed with PBS and lysed in plate (on ice) by adding 1ml ice cold lysis buffer (10mM Tris/HCl PH=7.5, 150mM NaCl, 0.5mM EDTA, 0.5% NP40,1uM PMSF). Cell lysis was transferred to individual tubes and homogenized on ice by passing through a 21-Gauge needle 10 times and cleared cell lysate was transferred to a new tube after centrifugation. An aliquot of 50 µl was set aside as “input”. Remaining lysate was incubated with 20 µl of anti-HA Affinity Gel (Sigma, EZview Red Anti-HA Affinity Gel, E6779) with rotation for 1 hour to overnight at $4^{o}$C. Samples were centrifuged and 50 µl of the supernatant was collected as "depleted supernatant". The the pulled down pellet (Co-IP) was washed with 750uL of lysis buffer for 3 times. Finally, loading buffer (2x Laemmli: 4% SDS, 20% glycerol, 120mM Tris-HCl, pH=6.8, 0.02% w/v bromophenol blue) was added to all samples, heated to 95°C for 10 minutes, and the pull down samples was flash cooled on ice before western analyses. For each experiment, 0.5% of input, 0.5% of depleted supernatant, and 20% pulldown samples from the Co-IP experiment were loaded into 15% SDS-polyacrylamide gel and transferred onto 0.45 µm nitrocellulose membrane (GE, 10600016). Blots were incubated with either rabbit-anti-Flag (1:10,000 CST 2368S) or rabbit-anti-HA (1:10,000 CST 3724), and then in HRP conjugated goat-anti-rabbit antibody (1:5,000 Santa Cruz, SC-2004), and immune detection was performed using Millipore chemiluminescent HRP substrate (Millipore, #WBKLS0100). Imaging was performed using XRS+ ChemiDoc imaging system (BioRad, 1708265).

*Purification of Recombinant Cdk2ap1 Protein*

To disrupt binding to CDK2, the previously described three amino acid binding domain was mutated, Thr108Ala, Glu109Ala, Arg110Ala (Referred to as "MutTER" from here on). ORFs of mouse $Cdk2ap1^{CAN}$, $Cdk2ap1^{CAN\text{-}}MutTER$, $Cdk2ap1^{\Delta N(MT2B2)}$ or, $Cdk2ap1^{\Delta N(MT2B2)}$-*MutTER* were each cloned into the pET28a bacterial expression vector (EMD Biosciences, 69864). The vector backbone was modified so that the cloned ORF would be downstream of an N-terminal cassette (His-Tag (6x)-Maltose Binding Protein (MBP)-short linker–TEV Cleavage site-ORF). Proteins were purified as previously described(*21*). Briefly, plasmids were transformed into E.coli LOBSTR expression cells (Kerafast, EC1002). Starter culture of 200 mL of LB liquid broth was grown at 30°C in the presence of Ampicillin and Chloramphenicol overnight. The following day, culture was added to pre-warmed (37°C) glassware and additional 1.3L of LB growth media. When an optical density of 0.5 at 600nm was reached, the bacteria culture was chilled to 16°C. Expression of protein was induced by adding IPTG to 250 µM (GoldBio 12481C5) and cultured overnight at 16°C. Cells were spun down and lysed in 20 mL of lysis buffer A (50 mM HEPES pH=7.5, 50 mM NaCl, 1 mM PMSF, 1 mM EDTA, 5 mg/mL Lysozyme, 30% glycerol) per 1.5 L of culture. Sample was incubated at Room Temperature (25°C) for 15 minutes while rocking, then 10 mL of lysis buffer B (50 mM HEPES pH=7.5, 300 mM NaCl, 1.5 mM PMSF, 15 mM β-mercaptoethanol, 30 mM imidazole, 20% Glycerol) per 1.5 L culture was added. Sample were then sonicated (On-pulse 10s, off-pulse 50s, amplitude 60%) on ice until proper viscosity was reached. Lysates were then cleared by centrifugation. His-tagged proteins were isolated using NI-NTA agarose (Qiagen, 30210), washed three times (50 mM HEPES pH=7.5, 150 mM NaCl, 1 mM PMSF, 5 mM β-mercaptoethanol, 20 mM imidazole, 20% glycerol), eluted with 2.5 mL elution buffer (50 mM HEPES pH=7.5,150 mM NaCl,5 mM β-mercaptoethanol, 250 mM imidazole, 20% glycerol). Samples were dialyzed overnight at 4°C (Fisher, 6-4033) in dialysis buffer (40 mM HEPES pH 7.5, 150 mM NaCl, 5 mM β-mercaptoethanol, 10% glycerol). Samples were further purified by size separation column purification via AKTA Chromatography through a Superdex 200 column (Millipore, G117-5175-01) in degassed purification buffer (50 mM HEPES, pH 7.5, 150 mM NaCl, 5 mM β-mercaptoethanol, 10% glycerol) followed by concentration of protein-containing fractions by three cycles of concentration/dilution using Amicon Ultra centrifugal filter units (MWCO 3 kDa, Millipore, Z647993). Protein concentration was determined using Nanodrop 2000 (A280nm). Concentrated proteins were aliquoted, flash-frozen in liquid nitrogen, and stored at −80°C.

*CDK2 Kinase Assay*

The CDK2 kinase assay was performed according to manufacturer's instructions (Promega, CDK2/CyclinE1 Kinase Assay, V4489). Briefly, we combined 2 µl enzyme mix (4ng CDK2/CyclinE1) and 2 µl substrate mix (0.1µg/µL Histone H1 and 150µM ATP) with various previously diluted 1 µl concentrations of recombinant mouse *Cdk2ap1*$^{CAN}$, *Cdk2ap1*$^{CAN}$ *Mut*, *Cdk2ap1*$^{\Delta N(MT2B2)}$ or , *Cdk2ap1*$^{\Delta N(MT2B2)}$ *Mut* proteins in 5 µl reactions, and incubated at room temperature for 60 minutes. After incubation, 5µl of ADP-Glo luminescent reagent was added, followed by a 40 min incubation at room temperature. Then 10 µl of Kinase Luminescence Detection Reagent was added and incubated at room temperature for 30 minutes. Sample luminescence were individually measured using Promega GloMax 20/20 Luminometer.

**Animals and Ethics Statement**

As a matter of caution and compliance, all appropriate authorizations have been acquired from institutional and/or federal regulatory bodies prior to performing this protocol. All mouse use, including but not limited to housing, breeding, production, sample collection for genotyping, and euthanasia, is in accordance with the Animal Welfare Act, the AVMA Guidelines on Euthanasia and are in compliance with the ILAR *Guide for Care and Use of Laboratory Animals,* and the UC Berkeley Institutional Animal Care and Use Committee (IACUC) guidelines and policies. Our animal care and use protocol (AUP-2015-04-7485-1) has been reviewed and approved by our IACUC for this project.

**Supplementary Text**

#### Supplementary Figure S1

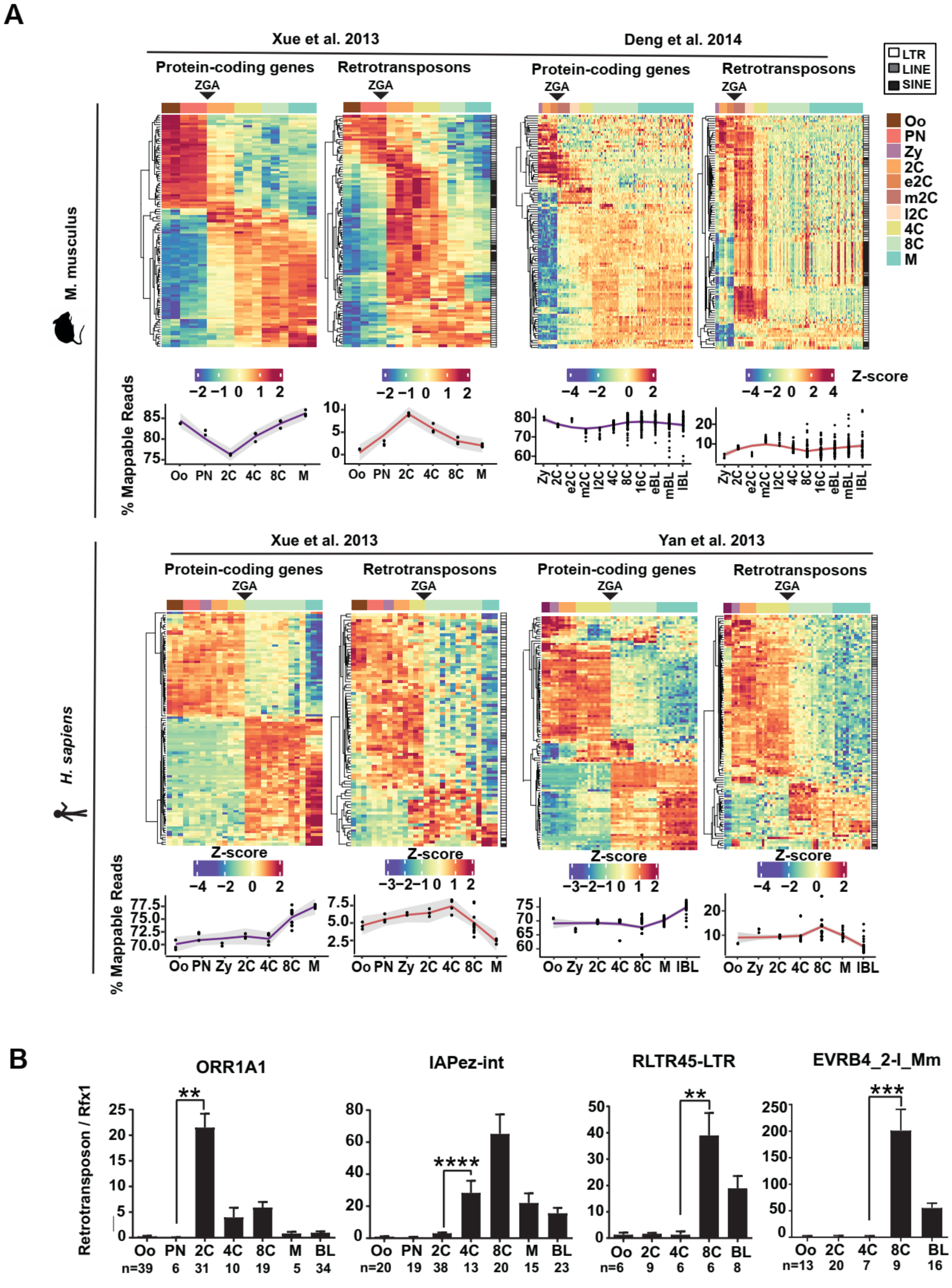

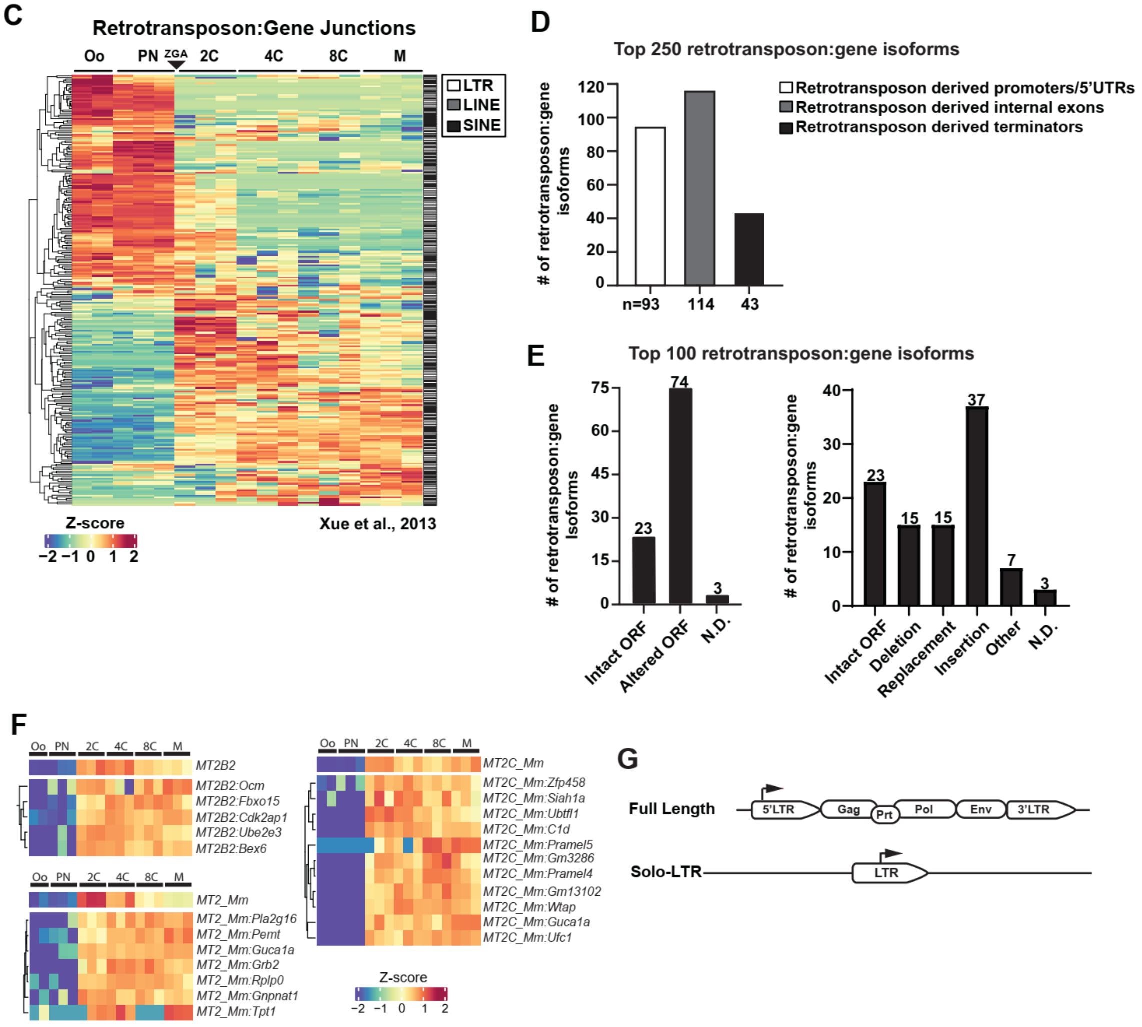
C
Retrotransposon:Gene Junctions
Oo
PN
ZGA
2C
4C
8C
M
LTR
LINE
SINE
Z-score
−2 −1 0 1 2
Xue et al., 2013
D
Top 250 retrotransposon:gene isoforms
### of retrotransposon:gene isoforms
120
100
80
60
40
20
0
n=93
114
43
Retrotransposon derived promoters/5'UTRs
Retrotransposon derived internal exons
Retrotransposon derived terminators
E
Top 100 retrotransposon:gene isoforms
### of retrotransposon:gene isoforms
75
60
45
30
15
0
74
23
3
Intact ORF
Altered ORF
N.D.
### of retrotransposon:gene isoforms
40
30
20
10
0
23
15
15
37
7
3
Intact ORF
Deletion
Replacement
Insertion
Other
N.D.
F
Oo PN 2C 4C 8C M
MT2B2
MT2B2:Ocm
MT2B2:Fbxo15
MT2B2:Cdk2ap1
MT2B2:Ube2e3
MT2B2:Bex6
MT2_Mm
MT2_Mm:Pla2g16
MT2_Mm:Pemt
MT2_Mm:Guca1a
MT2_Mm:Grb2
MT2_Mm:Rplp0
MT2_Mm:Gnpnat1
MT2_Mm:Tpt1
MT2C_Mm
MT2C_Mm:Zfp458
MT2C_Mm:Siah1a
MT2C_Mm:Ubtfl1
MT2C_Mm:C1d
MT2C_Mm:Pramel5
MT2C_Mm:Gm3286
MT2C_Mm:Pramel4
MT2C_Mm:Gm13102
MT2C_Mm:Wtap
MT2C_Mm:Guca1a
MT2C_Mm:Ufc1
Z-score
−2 −1 0 1 2
G
Full Length
5'LTR
Gag
Prt
Pol
Env
3'LTR
Solo-LTR
LTR

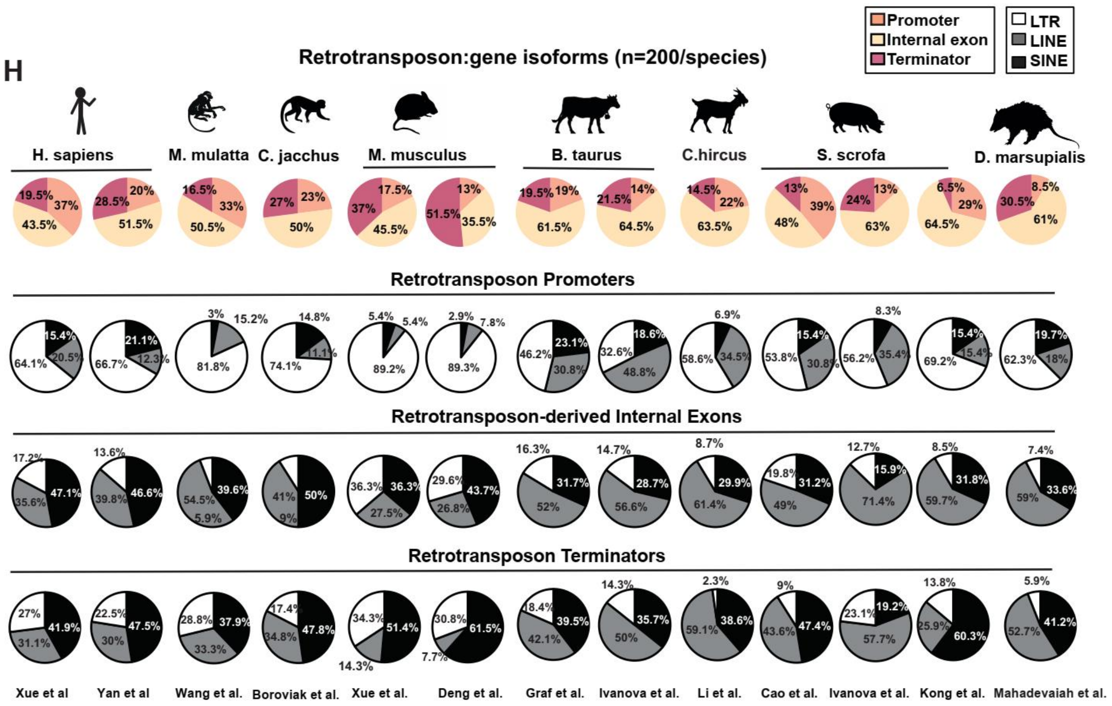
H
Retrotransposon:gene isoforms (n=200/species)
Promoter
Internal exon
Terminator
LTR
LINE
SINE
H. sapiens
M. mulatta
C. jacchus
M. musculus
B. taurus
C.hircus
S. scrofa
D. marsupialis
Retrotransposon Promoters
Retrotransposon-derived Internal Exons
Retrotransposon Terminators
Xue et al
Yan et al
Wang et al.
Boroviak et al.
Xue et al.
Deng et al.
Graf et al.
Ivanova et al.
Li et al.
Cao et al.
Ivanova et al.
Kong et al.
Mahadevaiah et al.

**Fig. S1.**

**Retrotransposons mediate gene regulation in mammalian preimplantation development.**
**A.** Retrotransposons and protein-coding genes exhibit similar preimplantation expression profiles in mice and human in multiple RNA-seq datasets. The top 100 most differentially expressed protein-coding genes and retrotransposon subfamilies are shown as heatmaps; the percentage of uniquely mapped reads that originate from protein-coding genes or retrotransposons are shown as line graphs. A subset of preimplantation stages are shown in heatmaps across all species to highlight the comparison among species. All the developmental stages that are available in the original datasets were included in line plots. White, LTR; grey, LINE; black, SINE; black triangle, ZGA; Z-score, the number of standard deviations from the expression mean of a protein-coding gene or a retrotransposon subfamily. Oo, oocyte; PN, pronucleus; 2C, two cell embryo; e2C, early two cell embryo; m2C, mid two cell embryo; l2C, late 2C embryo; 4C, four cell embryo; 8C, eight cell embryo; M, morula; BL, blastocysts; eBL, early blastocyst; mBL, mid blastocyst; lBL, late blastocyst; ZGA, zygotic genome activation. **B.** Single embryo real time PCR analyses confirm the dynamic expression of multiple retrotransposon subfamilies. Error bars, ± s.e.m., *P* values were calculated using unpaired, two-tailed Student's *t* test. ORR1A1, PN vs 2C, $^{**}P = 0.001$, t=3.5, df=35; IAPez-int, Oo vs PN, $^{****}P < 0.0001$, t=5.7, df=48; RLTR45-LTR, 4C vs 8C, $^{**}P = 0.001$ t=4.4, df=10; ERVB4-_2-I_Mm, 4C vs 8C, $^{***}P = 0.0005$, t=4.5, df=14. **C**. Retrotransposon-dependent gene isoforms were dynamically expressed in mouse preimplantation embryos. A heatmap shows the dynamic expression pattern of the top 250 most highly and differentially expressed splicing junctions between a retrotransposon and a proximal gene exon with a normalized retrotransposon-gene junction read counts $\geq 30$. **D.** The top 250 most highly and dynamically expressed retrotransposon:gene isoforms in mouse preimplantation embryos were broken down to three categories, including those that contain retrotransposon derived promoter/5'UTR, internal exon and terminator. **E.** Retrotransposon:gene isoforms frequently alter canonical ORFs. Manual curation prediction of ORFs from the top 100 most highly and dynamically expressed retrotransposon:gene isoforms in mouse preimplantation embryos. (Left) A bar plot shows the number of retrotransposon:gene isoforms with predicted ORFs that are either intact or altered. n.d., not determined. (Right) Those with predicted ORFs that differ from the canonical ORFs were further categorized as intact ORFs, deletion, replacement, insertion, other (those predicted with multiple N-terminal modification mechanisms) or N.D. (not determined). **F.** A diagram shows the gene structure of a full length LTR retrotransposon and a solo-LTR. **G**. Gene isoform driven by sequence related retrotransposon promoters exhibit an expression profile closely resemble that of the associated retrotransposon subfamilies. Representative heatmaps show the expression of a retrotransposon subfamily, including MT2_Mm, MT2B2 and MT2C_Mm and associated retrotransposon:gene isoforms across preimplantation stages. Z-score, the number of standard deviations from the expression mean of a protein-coding gene or a retrotransposon subfamily. Oo, oocyte; PN, pronucleus; 2C, two cell embryo; 4C, four cell embryo; 8C, eight cell embryo; M, morula. **H.** Retrotransposons provide alternative promoters, internal exons and terminators for gene isoforms in mammalian preimplantation embryos. The top 200 most dynamically expressed retrotransposon:gene isoforms in each mammalian species are analyzed for the LTR, SINE and LINE contribution to retrotransposon derived promoters, internal exons and terminators. RNA-seq data for these analyses were obtained from published datasets, documented in supplementary table S1-S3. All retrotransposon derived promoters contain predicted transcription start sites (TSSs) in our analyses.

#### Supplementary Figure S2

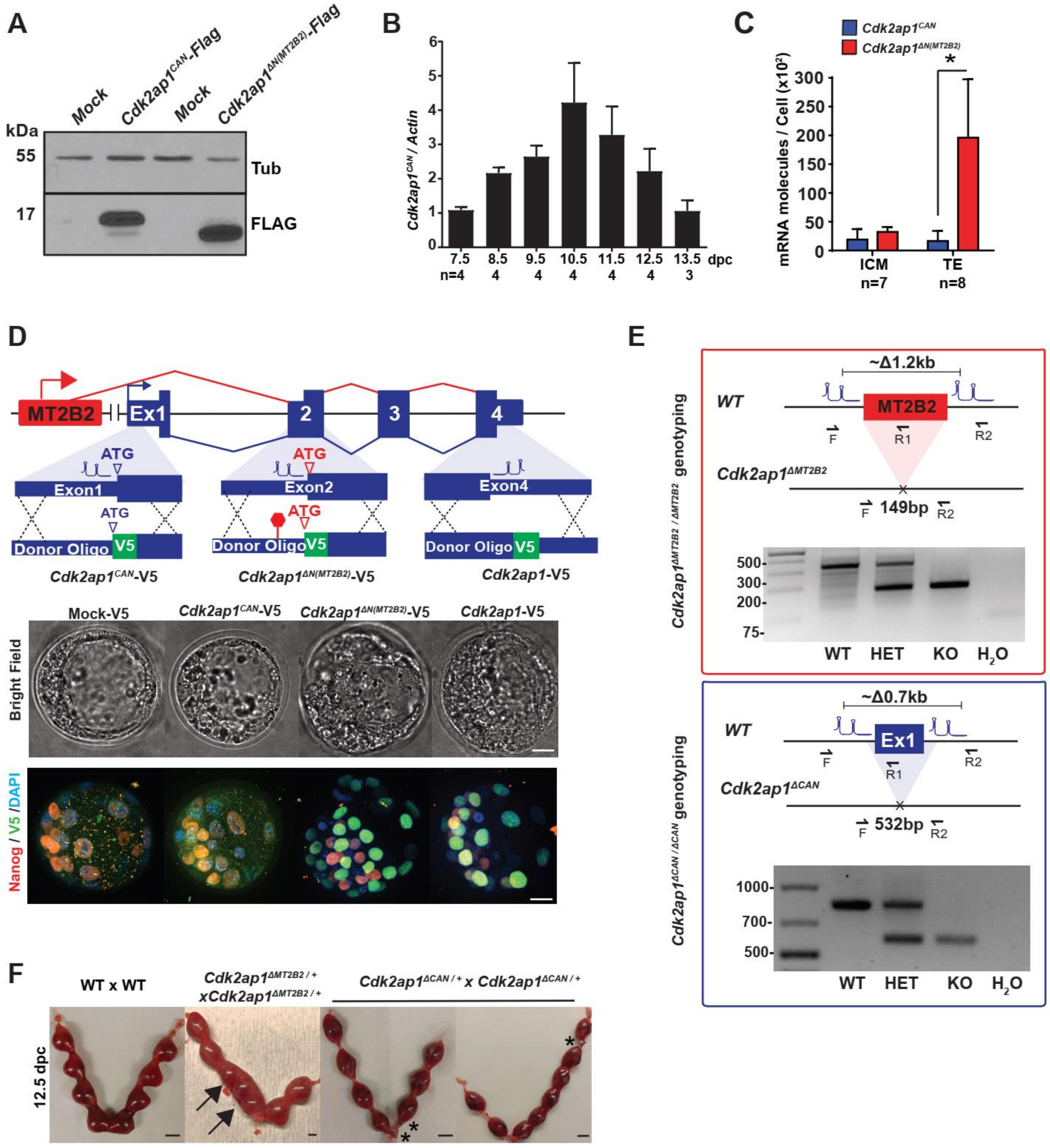

**Fig. S2.**

**A. Cdk2ap1$^{\Delta N(MT2B2)}$ and Cdk2ap1$^{CAN}$ exhibit distinct expression patterns.** **A.** The *Cdk2ap1*$^{\Delta N(MT2B2)}$ isoform produces a truncated Cdk2ap1 protein. Overexpression of *Cdk2ap1*$^{CAN}$ *-Flag* and *Cdk2ap1*$^{\Delta N(MT2B2)}$*-Flag* in HEK293T cells produce proteins of the expected sizes, demonstrating that the predicted ATG start codon in the *Cdk2ap1*$^{\Delta N(MT2B2)}$ isoform is functional for translation. . **B.** Real time PCR confirms the peak of *Cdk2ap1*$^{CAN}$ expression at 10.5 dpc embryos. **C.** *Cdk2ap1*$^{\Delta N(MT2B2)}$ expression is enriched in the TE of blastocysts. Absolute real time PCR quantitation of *Cdk2ap1*$^{CAN}$, *Cdk2ap1*$^{\Delta N(MT2B2)}$ and *Cdx2* was performed in isolated single cells of 4.0 dpc blastocysts. The expression of *Cdx2* was used to distinguish TE cells from ICM cells (Not shown). Error bars, s.e.m.; *Cdk2ap1*$^{CAN}$ vs. *Cdk2ap1*$^{\Delta N(MT2B2)}$ in TE, * $P$ =0.047, t=2.3, df=10. $P$ values are calculated using unpaired, two-tailed Student's $t$-test. **D.** Preimplantation specific Cdk2ap1 expression is mostly derived from the *Cdk2ap1*$^{\Delta N(MT2B2)}$ isoform. Using CRISPR-EZ, a V5 tag was engineered immediately after the canonical ATG for detection of Cdk2ap1$^{CAN}$ (*Cdk2ap1*$^{CAN}$*-Ex1-V5*). Additionally, a V5 tag was engineered immediately after the alternative ATG in exon 2 for detection of the Cdk2ap1$^{\Delta N(MT2B2)}$ isoform, and a stop codon (red hexagon) was introduced 12bp upstream of the Cdk2ap1$^{\Delta N(MT2B2)}$ ATG to prevent Cdk2ap1$^{CAN}$ production (*Cdk2ap1 –Ex2-V5*). Finally, a V5 tag was engineered immediately before the stop codon shared by all Cdk2ap1 isoforms for detection of total Cdk2ap1 (*Cdk2ap1-Ex4-V5*). Using immunostaining of V5, we detected no expression of Cdk2ap1$^{CAN}$-V5 in engineered blastocyst embryos, but a strong expression of Cdk2ap1$^{\Delta N(MT2B2)}$ and total Cdk2ap1 in TE. The V5 expression patterns of *Cdk2ap1 –Ex2-V5* and *Cdk2ap1-Ex4-V5* blastocysts are nearly identical, confirming that *Cdk2ap1*$^{\Delta N(MT2B2)}$ yields the majority of Cdk2ap1 proteins in preimplantation embryos. Two independent experiments were performed for each CRISPR-EZ editing experiment, with at least 5 embryos per condition. Scale bars, 20 µm. **E.** Diagrams illustrate the PCR genotyping strategies of CRISPR edited *Cdk2ap1*$^{\Delta MT2B2/\ \Delta MT2B2}$ (top) and *Cdk2ap1*$^{\Delta CAN/\Delta CAN}$ (bottom) embryos. Representative genotyping results are shown as electrophoresis images. **F.** Deletion of *Cdk2ap1*$^{\Delta N(MT2B2)}$, but not *Cdk2ap1*$^{CAN}$, is associated with embryo spacing defects during implantation. Uteri were collected at E12.5 from timed mating of wildtype x wildtype (n=40 uteri), *Cdk2ap1*$^{\Delta N(MT2B2)/+}$ x *Cdk2ap1*$^{\Delta N(MT2B2)/+}$ (n=34 uteri) and *Cdk2ap1*$^{\Delta CAN/+}$ x *Cdk2ap1*$^{\Delta CAN/+}$ (n=7 uteri) crosses. Embryo crowding is evident in uteri generated from *Cdk2ap1*$^{\Delta N(MT2B2)/+}$ x *Cdk2ap1*$^{\Delta N(MT2B2)/+}$ crosses 4. In comparison, resorption of correctly spaced embryos is observed in uteri from the *Cdk2ap1*$^{\Delta CAN/+}$ x *Cdk2ap1*$^{\Delta CAN/+}$ crosses (n=7), consistent with a post-implantation developmental defects speculated for *Cdk2ap1*$^{\Delta CAN/\Delta CAN}$ embryos. Black arrows, embryo crowding; *, resorbed embryos. Scale bars, 0.5 cm.

#### Figure S3

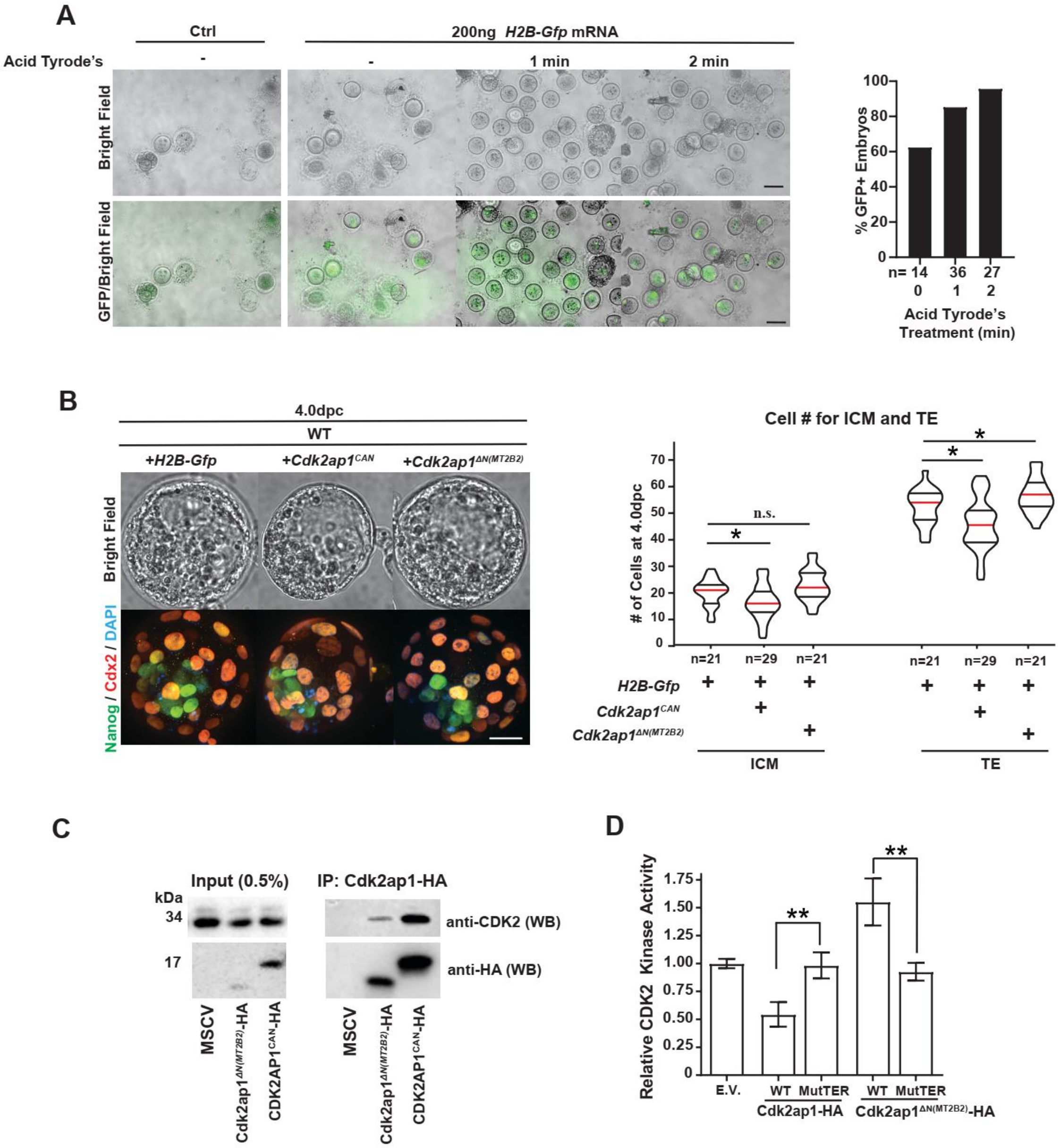
A
Ctrl
200ng H2B-Gfp mRNA
Acid Tyrode's
-
-
1 min
2 min
Bright Field
GFP/Bright Field
% GFP+ Embryos
n= 14
36
27
0
1
2
Acid Tyrode's Treatment (min)
B
4.0dpc
WT
+H2B-Gfp
+Cdk2ap1CAN
+Cdk2ap1ΔN(MT2B2)
Bright Field
Nanog / Cdx2 / DAPI
Cell # for ICM and TE
### of Cells at 4.0dpc
n.s.
n=21
n=29
n=21
H2B-Gfp
Cdk2ap1CAN
Cdk2ap1ΔN(MT2B2)
ICM
TE
C
Input (0.5%)
IP: Cdk2ap1-HA
kDa
34
17
anti-CDK2 (WB)
anti-HA (WB)
MSCV
Cdk2ap1ΔN(MT2B2)-HA
CDK2AP1CAN-HA
D
Relative CDK2 Kinase Activity
E.V.
WT
MutTER
Cdk2ap1-HA
Cdk2ap1ΔN(MT2B2)-HA

**Fig. S3.**

**$Cdk2ap1^{\Delta N(MT2B2)}$ and $Cdk2ap1^{CAN}$ have opposite functions in cell proliferation. A.** Acid Tyrode's treatment optimization to increase mRNA delivery efficiency into zygotes by electroporation. *H2b-Gfp* mRNA was electroporated into acid Tyrode's treated zygotes to determine the acid Tyrode's treatment duration for optimal zona pellucida thinning. Efficiency of mRNA delivery was measured by the percentage of GFP positive embryos 6 hours after electroporation. Representative GFP images (left) and quantification of GFP positive embryos (right) were shown. Two independent experiments were performed. **B.** *$Cdk2ap1^{CAN}$* and *$Cdk2ap1^{\Delta N(MT2B2)}$* have opposite effect on cell proliferation in wildtype preimplantation embryos. Wildtype embryos overexpressing *$Cdk2ap1^{CAN}$* or *$Cdk2ap1^{\Delta N(MT2B2)}$* were analyzed for ICM (Nanog+) and TE (Cdx2+) cell counts at 4.0 dpc using immunostaining (left). Quantitation was shown as violin plots with median (red line), lower (25%) and upper (75%) quartiles (black lines). *H2b-Gfp* vs *$Cdk2ap1^{CAN}$*: ICM, * $P = 0.03$, t=2.2, df=49; TE, *$P = 0.01$, t=2.7, df=49. *H2b-Gfp* vs *$Cdk2ap1^{\Delta N(MT2B2)}$*, ICM: n.s; TE, * $P = 0.03$, t=2.2, df=40. **C.** $Cdk2ap1^{CAN}$ and $Cdk2ap1^{\Delta N(MT2B2)}$ both bind endogenous mouse Cdk2. In HEK293T cells transfected with either C-Terminal HA-Tagged $Cdk2ap1^{CAN}$ ($Cdk2ap1^{CAN}$–HA) or C-Terminal HA-Tagged $Cdk2ap1^{\Delta N(MT2B2)}$ ($Cdk2ap1^{\Delta N(MT2B2)}$-HA), immunoprecipitation of HA pull down endogenous Cdk2. **D.** $Cdk2ap1^{CAN}$ and *$Cdk2ap1^{\Delta N(MT2B2)}$* have opposite effects on Cdk2 kinase activity. HEK293T cells overexpressing *$Cdk2ap1^{CAN}$-HA*, *$Cdk2ap1^{CAN}$-MutTER-HA*, *$Cdk2ap1^{\Delta N(MT2B2)}$-HA* or *$Cdk2ap1^{\Delta N(MT2B2)}$-MutTER-HA* were subjected to immunoprecipitation with anti-HA antibodies. Immunoprecipitated lysates were each incubated with recombinant CDK2, CYCLIN E, HISTONE H1 and ATP *in vitro*. Their effects on CDK2 activity were analyzed in a kinase assay. Error bars are means ± s.e.m. *$Cdk2ap1^{CAN}$* -HA vs *$Cdk2ap1^{CAN}$* -MutTER-HA, **$P = 0.009$, t=4.7, df=4; *$Cdk2ap1^{\Delta N(MT2B2)}$* vs *$Cdk2ap1^{\Delta N(MT2B2)}$*-MutTER-HA, **$P = 0.008$, t=4.8, df=4. Three independent experiments were performed. All *P* values were calculated based on the unpaired two-tailed Student's *t* test. n.s., not significant.

#### Supp Figure 4

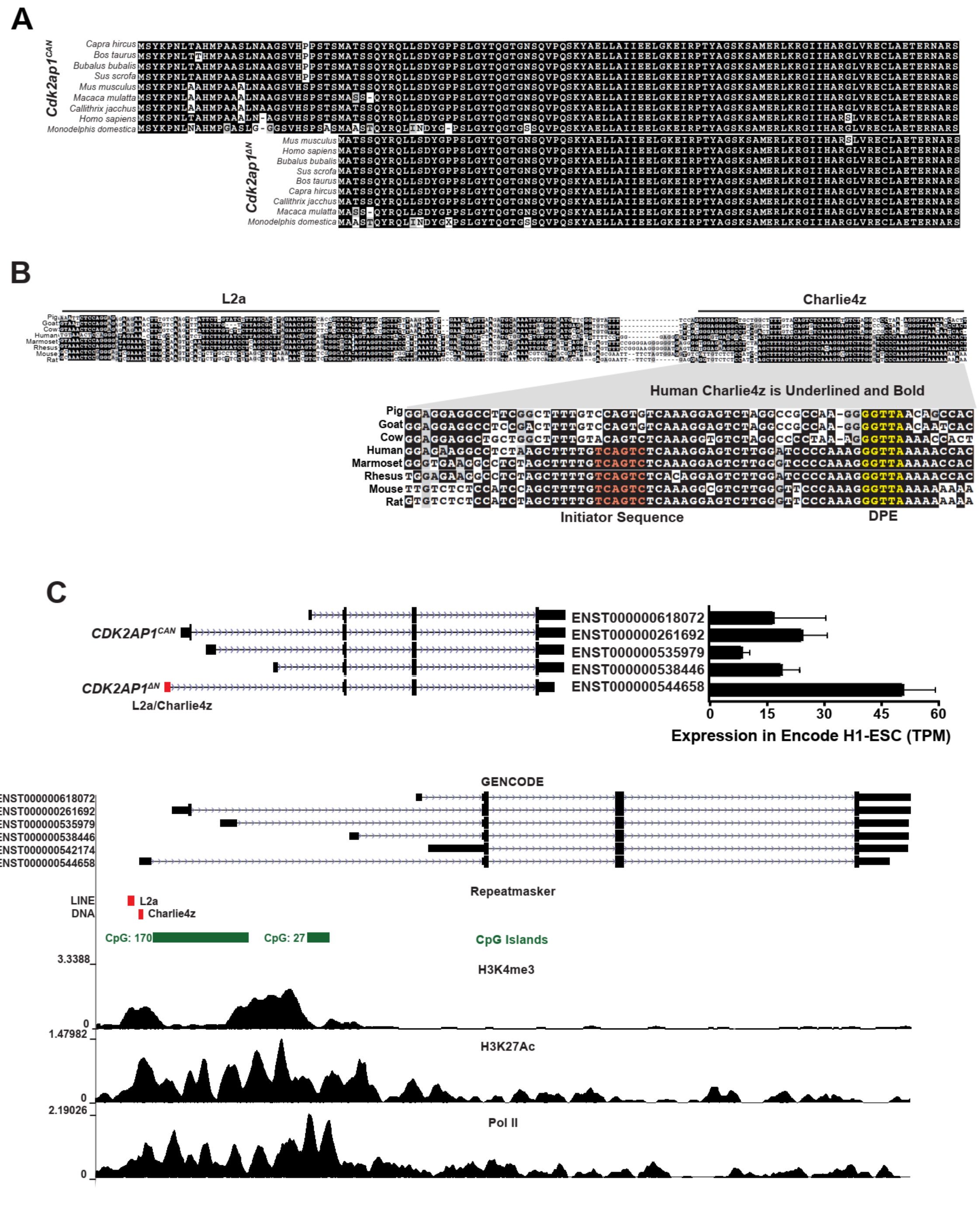

D

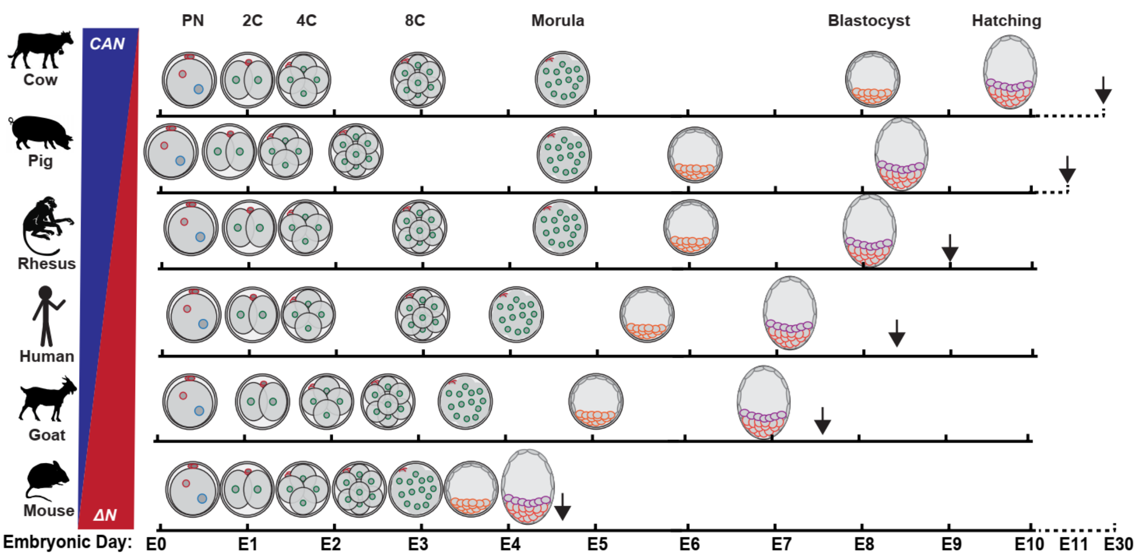

PN
2C
4C
8C
Morula
Blastocyst
Hatching
CAN
ΔN
Cow
Pig
Rhesus
Human
Goat
Mouse
Embryonic Day:
E0
E1
E2
E3
E4
E5
E6
E7
E8
E9
E10
E11
E30


E

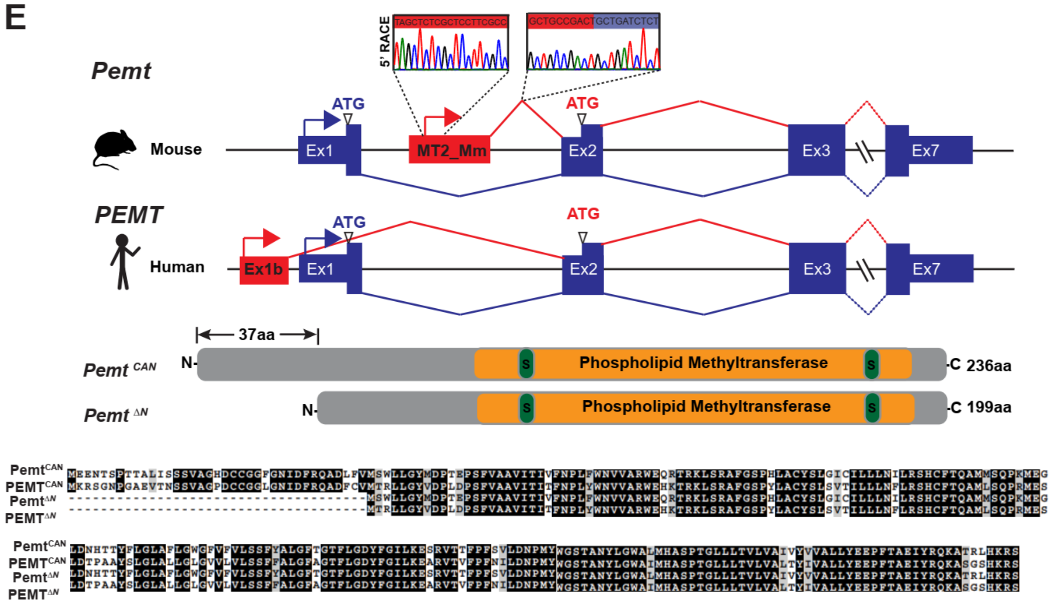

5' RACE
Pemt
Mouse
ATG
MT2_Mm
ATG
Ex1
Ex2
Ex3
Ex7
PEMT
Human
ATG
Ex1b
Ex1
ATG
Ex2
Ex3
Ex7
37aa
Pemt CAN
N-
S
Phospholipid Methyltransferase
S
-C 236aa
Pemt ΔN
N-
S
Phospholipid Methyltransferase
S
-C 199aa
PemtCAN
PEMTCAN
PemtΔN
PEMTΔN
PemtCAN
PEMTCAN
PemtΔN
PEMTΔN

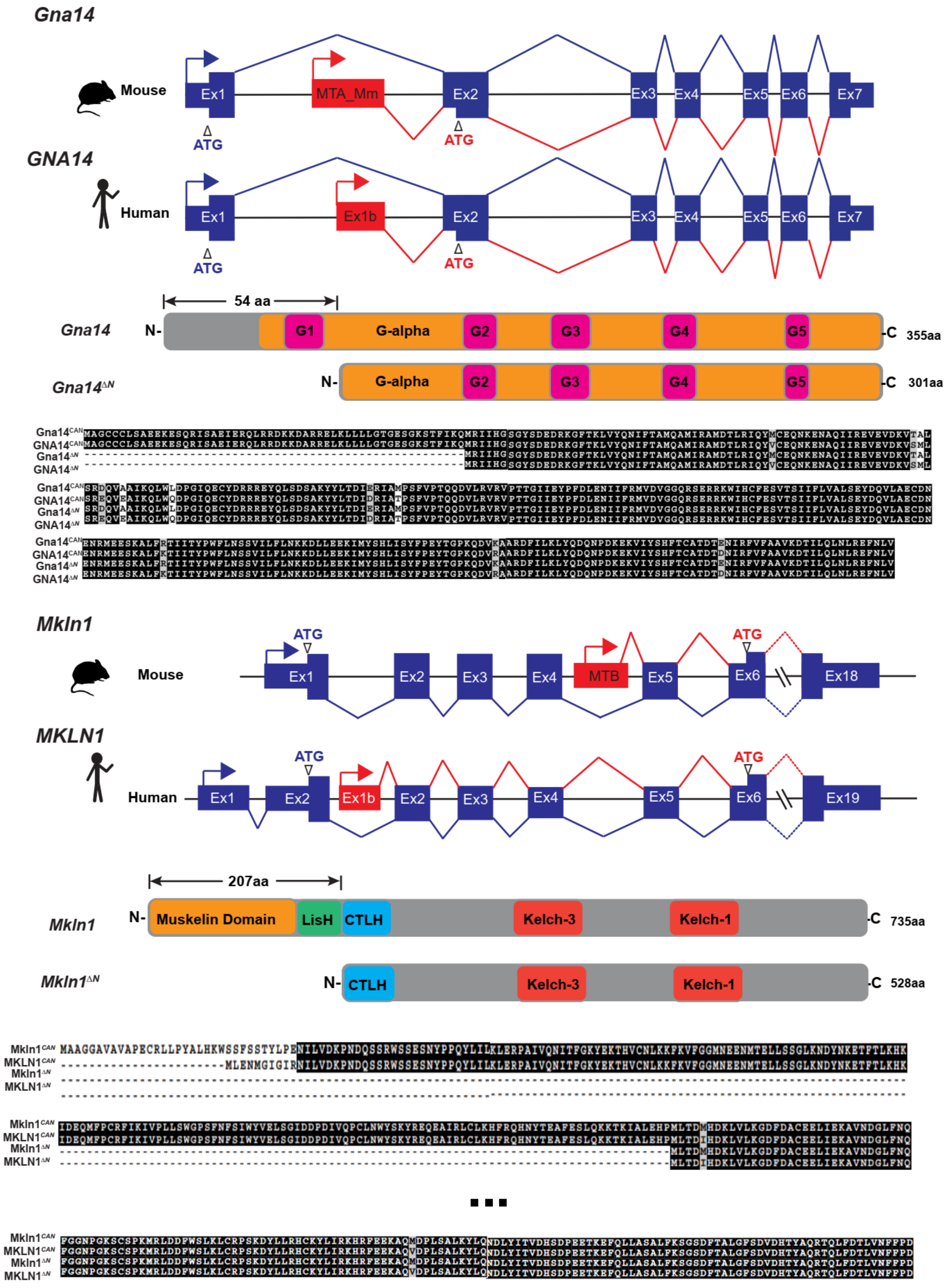

Gna14
Mouse
Ex1
MTA_Mm
Ex2
Ex3
Ex4
Ex5
Ex6
Ex7
ATG
ATG
GNA14
Human
Ex1
Ex1b
Ex2
Ex3
Ex4
Ex5
Ex6
Ex7
ATG
ATG
54 aa
Gna14
N-
G1
G-alpha
G2
G3
G4
G5
-C
355aa
Gna14ΔN
N-
G-alpha
G2
G3
G4
G5
-C
301aa
Mkln1
Mouse
ATG
ATG
Ex1
Ex2
Ex3
Ex4
MTB
Ex5
Ex6
Ex18
MKLN1
Human
ATG
ATG
Ex1
Ex2
Ex1b
Ex2
Ex3
Ex4
Ex5
Ex6
Ex19
207aa
Mkln1
N-
Muskelin Domain
LisH
CTLH
Kelch-3
Kelch-1
-C
735aa
Mkln1ΔN
N-
CTLH
Kelch-3
Kelch-1
-C
528aa

**Fig. S4.**

**The N-terminally truncated mouse Cdk2ap1$^{\Delta N(MT2B2)}$ protein is evolutionarily conserved in sequence and function. A.** Sequence alignment of *Cdk2ap1$^{CAN}$* and Cdk2ap1$^{\Delta N}$ isoforms across 8 mammalian species reveals a strong evolutionary conservation in protein sequences. **B.** In 8 eutherian mammals examined, alignment of the homologous genomic regions containing the L2a and Charlie4z fragments show sequence conservation. The region between L2a and Charlie4z is the least conserved, with goat, pig and cattle harboring a small deletion, and rodents and primates exhibiting sequence variations. The Charlie4z element contains a predicted initiator sequence (red) and a DPE (Downstream Promoter Element, yellow), both of which implicate promoter functionality. **C.** Chip-seq and RNA-Seq strongly suggest the L2a/Charlie4z is a bona fide promoter in human ES Cells. (Top) Each of the five human isoforms for CDK2AP1 were quantified by RNA-seq for expression in H1 embryonic stem cells from ENCODE project. (Bottom) Signatures of active promoter (H3K4me3, H3K27Ac, and Pol II) in human embryonic stem cells were illustrated with ChIP-seq data from ENCODE and Roadmap Epigenomics project. (GSE23316)(*1*) were downloaded from NCBI GEO database(*2*), and Kallisto(*3*) was used to quantify the isoform expression levels with GENCODE (GRCh38 ver. 26)(*4*) . **D.** A diagram illustrates the approximate durations of preimplantation development in different mammalian species. Arrow, approximate timing of implantation. **E.** Multiple mouse retrotransposon promoters yield specific gene isoforms that encode N-terminally altered ORFs orthologous to Refseq and/or Ensemble annotated human gene isoforms. Gene structure, protein motif analyses and sequence alignment were shown for *Pemt*, *Gan14* and *Mkln1*. For *Pemt*, the transcriptional start site, as well as the splicing between retrotransposon and gene exon, were experimentally validated by 5' RACE and RT-PCR.

Tables S1, S2 and S3 exceed size limit and can be found on third-party host (https://www.dropbox.com/sh/auo40kyfgu5jaiz/AABGEs2pR_EkHl3Pkgwldv4ya?dl=0)

**Table S1.**

RNA-seq analyses of retrotransposon expression in mouse, primate and livestock preimplantation embryos.

**Table S2.**

RNA-seq analyses of protein-coding gene expression in mouse, primate and livestock preimplantation embryos

**Table S3.**

RNA-seq analyses of retrotransposon:gene junction reads in mouse, primate and livestock preimplantation embryos.

**Table S4.**

Experimental validation of selected retrotransposon:gene isoforms.

**Table S5.**

ORF analyses on retrotransposon:gene isoforms in preimplantation embryos by manual curation.

**Table S6.**

RNA-seq analyses of Cdk2ap1 gene isoforms in mouse, primate and livestock preimplantation embryos.

**Table S7.**

An overview of the timing of preimplantation development in different mammals

**Table S8.**

Human-mouse conservation analysis of retrotransposon-gene isoforms with ORF alterations

**Table S9.**

Primer/oligo design for genotyping, CRISPR editing, real time PCR and RACE.
